# NRF2 translation block by inhibition of cap-dependent initiation sensitizes lymphoma cells to ferroptosis and CAR-T immunotherapy

**DOI:** 10.1101/2024.09.09.612133

**Authors:** Paola Manara, Austin D. Newsam, Venu V. G. Saralamma, Marco V. Russo, Alicia Bilbao Martinez, Nikolai Fattakhov, Tyler A. Cunnigham, Abdessamad Y. Alaoui, Dhanvantri Chahar, Alexandra M. Carbone, Olivia B. Lightfuss, Alexa M. Barroso, Kyle Hoffman, Francesco Maura, Daniel Bilbao, Jonathan H. Schatz

**Affiliations:** University of Miami Miller School of Medicine Sheila and David Fuente Graduate Program in Cancer Biology, University of Miami Miller School of Medicine, Miami, FL, US; University of Miami Miller School of Medicine Medical Scientist Training Program, Miami, Fl, US; University of Miami Miller School of Medicine Molecular and Cellular Pharmacology Graduate Program, Miami, FL, US; Division of Hematology, Department of Medicine, University of Miami Miller School of Medicine, Miami, Fl, US; Sylvester Comprehensive Cancer Center, Miami, FL, US; Department of Pathology and Laboratory Medicine, University of Miami Miller School of Medicine, Miami, Florida, US; Bioinformatics Solutions Inc., Waterloo, ON N2L 6J2, Canada; Division of Myeloma, Department of Medicine, University of Miami Miller School of Medicine, Miami, Fl, US

**Author notes:** Conflict of interest statement Disclosure: The authors declare no potential conflicts of interest.

## Abstract

Cancers coopt stress-response pathways to drive oncogenesis, dodge immune surveillance, and resist cytotoxic therapies. Several of these provide protection from ferroptosis, iron-mediated oxidative cell death. Here, we found dramatic sensitization to ferroptosis upon disruption of cap-dependent translation in diffuse large B-cell lymphoma (DLBCL). Specifically, rocaglate inhibitors of the eIF4A1 RNA helicase synergized with pharmacologic ferroptosis inducers, driven by a collapse of glutathione production that protects polyunsaturated fatty acids from ferroptotic oxidation. These effects occur despite initial up-regulation of specific protective factors. We find lost translation of NRF2, oncogenic master regulator of antioxidant gene-expression, is a key consequence of eIF4A1 inhibition. In vivo, combination of the clinical rocaglate zotatifin with a pharmacologically optimized ferroptosis inducer eradicated DLBCL patient derived xenografts. Moreover, we found zotatifin pre-exposure sensitized DLBCL to CD19-directed chimeric antigen receptor (CAR-19) T cells. Translational disruption therefore provides new opportunities to leverage therapeutic impacts of ferroptosis inducers including cytotoxic immunotherapies.

**Significance:** We find translational disruption sensitizes lymphomas to ferroptosis, enhancing efficacy of CAR-T cells and multiple drugs. NRF2 loss mediates these effects, informing promising new therapeutic combinations. Multiple cancers exploit NRF2 to resist a wide variety treatments. These results expand therapeutic implications from its loss downstream of eIF4A1 inhibition.

## Introduction

Ferroptosis is iron-mediated cell death triggered by compounds that reduce intracellular glutathione (GSH) or inhibit other mechanisms of protection from membrane lipid oxidation (1,2). Many cancer cells resistant to other treatments retain sensitivity to pharmacologic inducers of ferroptosis since expanded labile iron pools facilitate malignant proliferation. Clinical trials to date, however, have failed to establish clear therapeutic window for ferroptosis inducers in cancer patients.

Previous work shows cells with reduced levels of protein synthesis like hematopoietic stems cells (HSCs) have increased sensitivity to ferroptosis due to reduced presence of protective pathways (3). Notably, activation of the antioxidant-response transcription factor NRF2 (nuclear factor erythroid 2-related factor 2), promotes resistance to rocaglate cap-dependent translation inhibitors, evidenced in a CRISPR/Cas9 screen in cultured murine pro-B cells (4). The mechanism by which NRF2 mediates rocaglate resistance, however, remains elusive. We previously showed these drugs trigger complex and dynamic protein expression changes suggesting translational inhibition causes a stress response that drives some proteins to increase in expression even as many key oncogenic factors decline (5). Among other things, we found increased production during rocaglate exposure of SLC3A2 also known as CD98 heavy chain that together with SLC7A11 (xCT) comprises system X_c_^—^. This heterodimeric amino-acid exchanger plays a key role in ferroptosis protection by importing cystine to fuel GSH production by the cytoplasmic enzyme GPX4 (6). Rocaglates targeting the eIF4A translation-initiation factor include the clinical compound zotatifin (eFT226), which advanced to clinical trials for solid tumors (7,8). Interplay between ferroptosis and translation inhibition remains unexplored.

B-cell malignancies like diffuse large B-cell lymphoma (DLBCL) are driven by deregulation of several oncoproteins that rely on cap-dependent translation mediated by eIF4F, the mRNA cap-binding complex that includes eIF4A as its enzymatic core (9,10). Targeting eIF4F bypasses drug-resistance driven by redundant signaling pathways (11). With up to 40% of DLBCL patients experiencing relapsed or refractory (R/R) disease and facing poor outcomes, the advent of immunotherapy, particularly chimeric antigen receptor T cells targeting CD19 (CAR-19), has transformed the treatment landscape, curing up to 30-40% of individuals with relapsed/refractory (rel/ref) DLBCL (12–15). Nonetheless, the majority of rel/ref patients – especially considering high expense and production times that limit CAR-19 availability – ultimately will die as a result of lymphoma progression. Therefore, novel therapeutic approaches are urgently needed.

In this study, we assessed zotatifin’s impact on transcription and translation in DLBCL cells. Transcriptionally activated stress response factors failed to be effectively translated to their protein products. Translational upregulation of ferroptosis-protective proteins surprisingly did not provide protection from ferroptosis inducers. To the contrary, zotatifin demonstrated pronounced synergy with a wide variety of compounds that induce ferroptosis across multiple cellular systems in vitro and against DLBCL patient-derived xenograft (PDX) tumors in vivo. Synergy extended also to zotatifin plus INFγ, a cytotoxic T-cell effector cytokine that induces ferroptosis in target cells (16). We therefore assessed CAR-19 therapies and found pre-treatment with zotatifin significantly enhanced sensitivity.

## Results

### Rocaglates trigger transcriptional stress responses that are blocked translationally

Disruption of cap-dependent initiation by rocaglates reprograms translation, and we previously established conditions to examine reprogramming by mass spectrometry (5). Because B-cell lymphomas are among the most promising tumor types for rocaglate therapy (9–11), we applied these techniques to DLBCL, employing SU-DHL10 cells derived from the germinal center B-cell (GCB) subtype of DLBCL. In parallel, we performed RNA sequencing (RNA-seq) to examine relationships between mRNA and protein expression alterations. RNA-seq analysis revealed differential expression of 4450 genes, 2749 upregulated and 1703 downregulated, in response to zotatifin (**Fig. S1A**). Gene-set enrichment analysis (GSEA) revealed expression changes in seven HALLMARK gene sets, 4 up-regulated, 3 down-regulated at nominal p<0.01 and FDR <0.25 (**Fig. S1B**). We observed upregulation of stress responses promoting survival, including ‘TNFα Signaling via NF-κB’ and ‘Hypoxia’ (**Fig. S1C**). Further studies validated these findings, including induction of p65 phosphorylation and activation of an eGFP NF-kB reporter (**Fig. S1D-F**). The most downregulated pathways included ‘E2F Targets’, consistent with the well-described growth arrest in response to rocaglates (**Fig. S1G**) (5,17). Bulk transcriptomics analysis therefore revealed activation of stress-responsive mRNA expression programs after rocaglate treatment.

Under treatment with a translation inhibitor, transcriptionally up-regulated genes may not efficiently lead to increased protein products. We therefore assessed the translatome by TMT-pSILAC and mass-spectrometry (MS) (**Fig. 1A**). As previously described (5), we employed conditions to examine translation during the final 8 hours of a 24-hour treatment window using rocaglate concentrations that disrupt translation without significant cell death (**Fig. S2A**). We designed these conditions to capture cellular adaptation to the well-described initial rapid decline in key survival factors to find proteins that increase despite eIF4A disruption. Zotatifin up-regulated 88 proteins (log_2_FC>1, p<0.01), including the previously reported SLC3A2 (**Fig. 1B**) (5). Down-regulated proteins under these conditions (**Fig. S2B**) reflected the effects of lost key primary targets as shown also from HALLMARK GSEA analysis including MYC, the mTOR pathway, G2M-Checkpoints, and E2F targets (**Fig. S2C)** (10,17,18).

**Figure 1:**
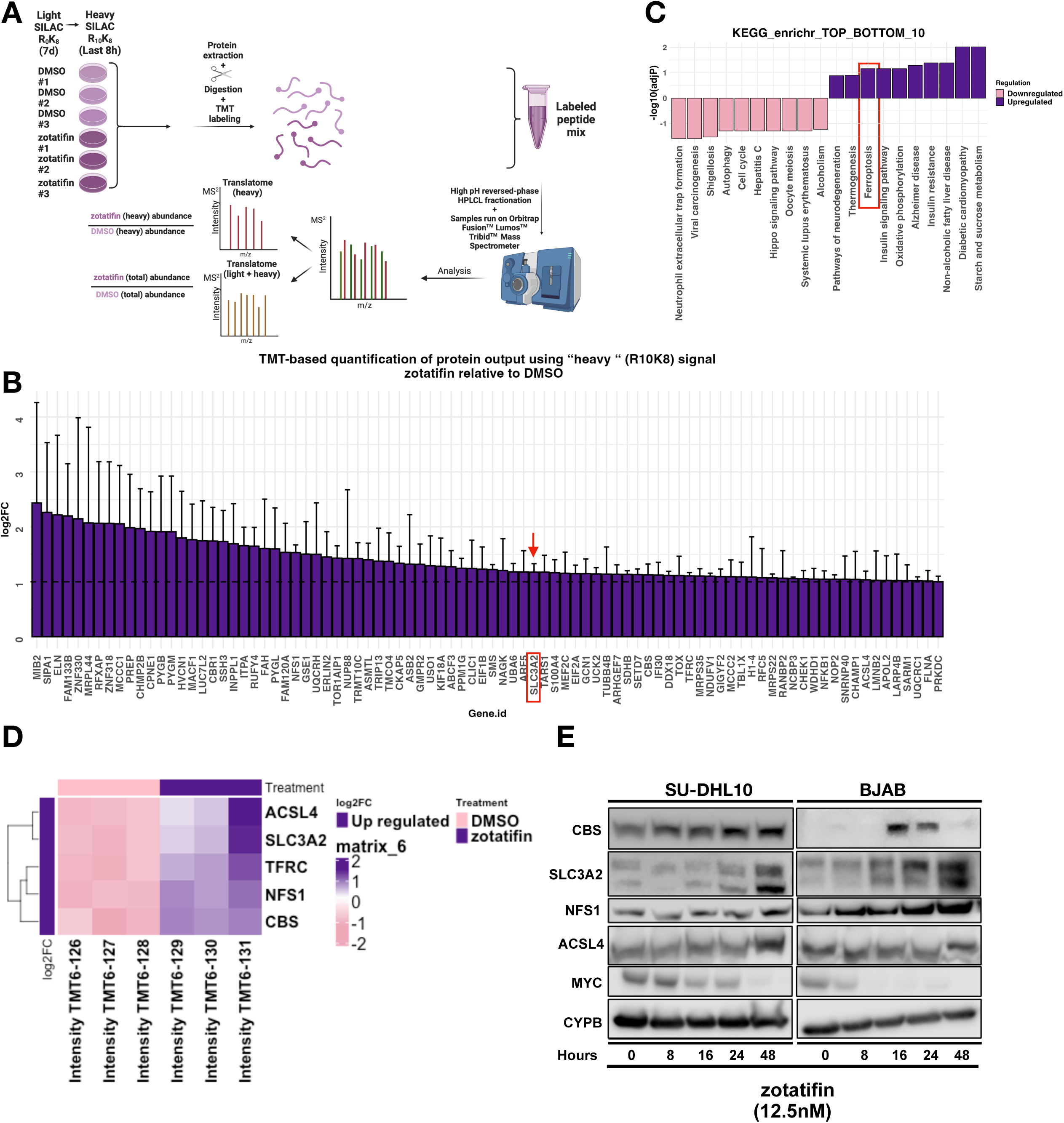
A) TMT-pSILAC workflow used to identify zotatifin-induced changes in protein output in DHL10 cells. B) TMT-pSILAC analysis showing upregulated proteins with a log_2_FC>1. C) Enrichr KEGG pathway analysis of significantly overrepresented genes (log_2_FC ≤-1 and log_2_FC ≥1). The 10 most significant pathways are displayed based on -log10(adjusted P-value) (−log_10_(adjP)≤−1 and ≥1 -log_10_(adjP)) in KEGG pathway analysis. D) Heatmap showing ferroptosis-related genes identified in the TMT-pSILAC dataset with a log_2_FC ≥1. E) Immunoblot time course analysis of SU-DHL10 (left) and BJAB (right) cells treated with 12.5 nM zotatifin for the indicated durations.

Strikingly, comparing TMT-pSILAC significantly upregulated proteins and corresponding mRNA transcripts, we found only a single gene, *NOP2,* enconding nucleolar protein 2, was significantly upregulated in both data sets (**Fig. S2D-E**). Transcriptional stress responses activated by rocaglates are thus hampered from being effectively translated into functional proteins, revealing a decoupling of mRNA and protein expression. Moving forward, we focused exclusively on protein alterations as the key functional output.

### Multiple ferroptosis protection proteins are translationally regulated in response to rocaglates

KEGG enrichment analysis of up-regulated proteins revealed several pathways with no connection to lymphoma or B-lymphocyte biology, suggesting many up-regulated proteins are not coordinately rescued along biologic ontologies (**Fig. 1C**). An exception was “Ferroptosis,” to which B-cell lymphomas are specifically susceptible (19,20). Up-regulated proteins included five ferroptosis mediators, four suppressors: cysteine desulfurase (NFS1) (21), cystathionine beta-synthase (CBS) (22), SLC3A2 (23) and TFR1 (24), plus the ferroptosis facilitator ACSL4 (25) (**Fig. 1D**). Immunoblotting confirmed upregulation of these factors during zotatifin exposure (**Fig. 1E**). These findings suggest a reshaping of the translatome toward protection against ferroptosis, though with unclear impact on drug efficacy since induction of ferroptosis is not a known consequence of rocaglate treatment. Hence, we examined gene and protein expression data from the Cancer Cell Line Encyclopedia (CCLE) (26), enabling cross-comparisons with drug-sensitivity data available on DepMap. This dataset includes sensitivity information regarding the synthetic rocaglate CR-1-31B (Table 1). We plotted CR-1-31B EC_50_ values against mRNA expression of *SLC3A2, NFS1, CBS, ACSL4, and TFRC*, genes encoding the proteins upregulated in our TMT-pSILAC analysis (**Fig. 1D-E**), as well as the ferroptosis-protective genes *GPX4* and *FTH1*. Decreased sensitivity to CR-1-31B associates with increased expression of *SLC3A2, NFS1, GPX4,* and *FTH1* mRNA, while slopes for *ACSL4*, *TFRC,* and *CBS* were not statistically significant (**Fig. 2A**). For functional validation, we engineered inducible SLC3A2 overexpression in SU-DHL-10 cells and confirmed decreased rocaglate sensitivity (**Fig. 2B-D**). In addition, we obtained a previously engineered *SLC3A2* knockout clone of U266 multiple myeloma cells (27), which showed increased zotatifin sensitivity (**Fig. 2E**). These findings suggest an induced protective response against ferroptotic stress results from rocaglate treatment, prompting further exploration ferroptosis’ role in rocaglates’ anti-lymphoma potency.

**Figure 2:**
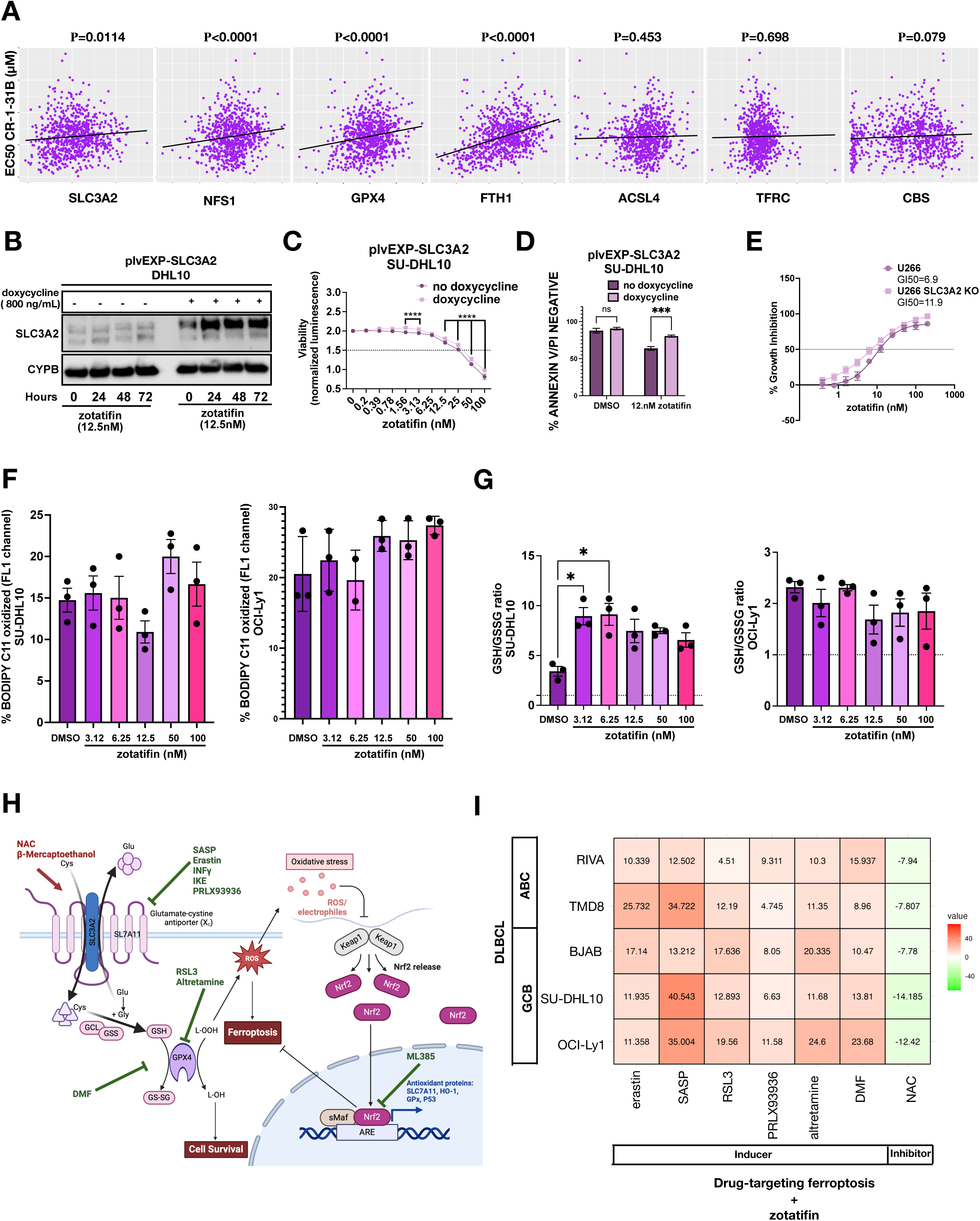
A) Linear regression slope analysis of Cancer Cell Line Encyclopedia (CCLE) data for mRNA expression of *SLC3A2, NFS1, GPX4, FTH1, ACSL4, TFRC*, and *CBS* versus CR-1-31B EC_50_, generated using the Dependency Map (DepMap) portal. B) Immunoblot analysis of SU-DHL10 cells engineered with doxycycline (doxy)-inducible SLC3A2 expression versus non-induced cells, treated with 12.5 nM zotatifin for 24, 48, and 72 hours. Data from 2 independent experiments. C-D) Effects of SLC3A2 overexpression in SU-DHL10 cells treated with 12.5 nM zotatifin for 48 hours, as measured by (C) ATP-dependent viability assay and (D) Annexin-V/PI staining. C) and D) Statistical analysis was performed using repeated measures two-way ANOVA with Sidak’s multiple comparison test ***P < 0.001, ****P < 0.0001). E) Growth inhibition assay of U266 MM cell lines with or without CRISPR-mediated SLC3A2 knockout. F) Measurement of oxidized BODIPY C11 fluorescence via flow cytometry across varying concentrations of zotatifin (6.5 nM to 100 nM) in SU-DHL10 (left), OCI-Ly1 (right). G) GSH/GSSG ratio calculated using the concentrations of GSH and GSSG at different zotatifin concentrations (6.5 nM to 100 nM) in SU-DHL10 (left) and OCI-Ly1 (right) cells. F) and H) Statistical analysis was performed using one-way ANOVA with Bonferroni’s multiple comparison test. Data represent the mean ± SEM of three independent experiments (*P < 0.05 vs. DMSO). H) Schematic representation of molecular ferroptosis protection mechanisms and targeted pathways of ferroptosis. I) Bliss δ synergy score heatmap for zotatifin in combination with various ferroptosis inducers (erastin, SASP, RSL3, DMF, PRLX93936, Altretamine) and inhibitor (NAC) in RIVA, TMD8, BJAB, SU-DHL10, and OCI-Ly1 DLBCL cell lines. A Bliss δ synergy score > 10 indicates synergy, between −10 and 10 suggests an additive effect, and < −10 indicates antagonism.

**Table 1:** Potent activity of rocaglates against aggressive B-lymphoma cell lines. The table includes viability data for the synthetic rocaglate CR-1-31B sourced from the Cancer Cell Line Encyclopedia (CCLE) with assessments performed at 96 hours. Our results for rocaglamide A (rocA), silvestrol, and zotatifin we performed at 48 hours.

| Cell Line | CR-1-31B* | RocA | Silvestrol | Zotatifin |
| --- | --- | --- | --- | --- |
| OCI-Ly3 | 0.96 | 1.60 | 2.10 | 4.30 |
| TOLEDO | 2.29 | 2.10 | 4.80 | 6.50 |
| SU-DHL-4 | 3.48 | 1.70 | 2.20 | 7.30 |
| KARPAS422 | 3.71 | 0.70 | 1.10 | 2.20 |
| OCI-Ly19 | 3.88 | 1.30 | 2.40 | 4.20 |
| SU-DHL-10 | 6.75 | 3.00 | 5.20 | 13.20 |
| BJAB |  | 1.5 | 2.6 | 10.8 |
| SU-DHL-6 |  | 2.30 | 3.90 | 8.10 |

\*96-hr data from CCLE; other compounds 48 hrs

### Rocaglate-treated cells show profound sensitivity to induction of ferroptosis

BODIPY 581/591 C11 detects lipid peroxide levels during onset of ferroptosis (28). We assessed BODIPY by flow cytometry after treatment with zotatifin (**Fig. 2F**). Interestingly, while zotatifin caused little or weak evidence of ferroptosis, there were significant cellular increases in the anti-ferroptotic metabolite glutathione (GSH) compared to its oxidized form glutathione disulfide (GSSG) in SU-DHL10 but not in OCI-Ly1 (**Fig. 2G**). This suggest an induction of ferroptotic stress by rocaglates that may be compensated by the translationally mediated up-regulation of protective factors and wanted to test if pharmacologic ferroptosis inducers would overcome this protection. System x_c_^−^ inhibitors (**Fig 2H**) led to the original discovery of ferroptotic cell death (1) and are under clinical evaluation in the treatment of cancer. We assessed rocaglates in combination with erastin, PRLX-93936, and sulfasalazine (SASP). We also tested RSL3 and dimethyl fumarate (DMF), which counteract anti-ferroptotic activities of GPX4, plus the ferroptosis-inducing chemotherapeutic altretamine. As a negative control, we employed the potent antioxidant and ferroptosis protective molecule N-acetyl cysteine (NAC). **Fig. 2I** summarizes results of synergy assessments in multiple lines using the Bliss-independence model, revealing potent synergy across multiple rocaglate-ferroptosis inducer combinations.

### Zotatifin enables oxidation-driven damage to membranes by ferroptosis inducers

We next investigated GSH/GSSG ratios that indicate cellular protection from ferroptosis (2). System x_c_− inhibitors fully counteracted the protective GSH increase induced by zotatifin in DHL-10 cells and led to a significant reduction in OCI-Ly1 cells (**Fig. 3A**). Previous studies report contrasting findings regarding activation of apoptotic markers during ferroptosis. While Dai et al. found activation upon ferroptosis induction (29), others (30) described a mechanism that is not strictly connected to apoptosis, although both processes can be triggered under certain stress conditions, such as endoplasmic reticulum stress (31). It’s known that rocaglate-induced translation inhibition triggers apoptosis by increasing caspase 3 activity (5), reducing MCL1 translation (11), and disrupting components of mitochondrial integrity (32). We assessed these events in response to zotatifin alone and in combination with ferroptosis inducers and observed activation of cleaved-caspase 3 and decreased levels of X-chromosome-linked inhibitor of apoptosis protein (XIAP), likely due to the combination of ferroptosis induction and zotatifin treatment (**Fig. 3B-C, Fig. S3B**). Consistent with previous observations, GPX4 declined in response to ferroptosis inducers but was not impacted by zotatifn alone. The decline in MYC expression is positive control for rocaglate-induced disruption of translation (10,11). These combinations appear to synergistically kill cancer cells through both apoptosis and ferroptosis. BODIPY staining showed a significant increase in lipid peroxidation when zotatifin was combined with erastin, RSL3, or DMF (**Fig. 3D**). Conversely, a consistent decrease in lipid peroxidation was observed with the antioxidant NAC. We also measured ROS levels using MitoSOX Red and DHE fluorescent probes, which detect superoxide anion and found significant increases when zotatifin was combined with the other agents (**Fig. 3E-F**). These findings reveal that rocaglates broadly synergize with pharmacologic ferroptosis inducers by promoting ROS production, causing polyunsaturated fatty acid (PUFA) plasma membrane damage leading to death.

**Figure 3:**
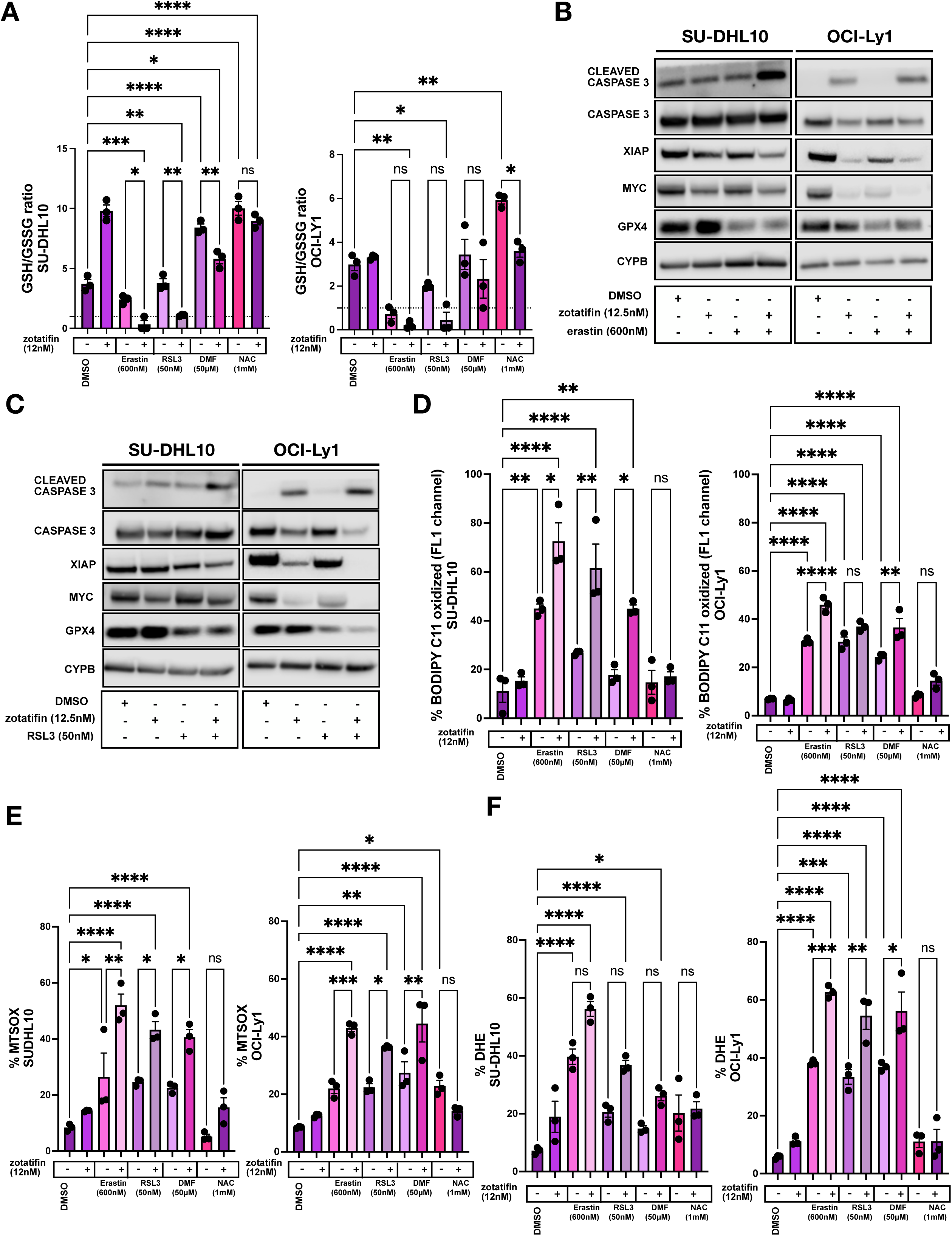
A) GSH/GSSG ratio in SU-DHL10 (left) and OCI-Ly1 (right) cells treated with zotatifin alone or with ferroptosis inducers (erastin, RSL3, DMF) or inhibitor (NAC). Statistical analysis: one-way ANOVA with Bonferroni multiple comparison test. Data represent mean ± SEM of 3 independent experiments. Statistical analysis used one-way ANOVA with Bonferroni multiple comparison test (*P < 0.05, **P < 0.01, ***P < 0.001, ****P < 0.0001). B-C) Immunoblot for cleaved caspase 3, XIAP, MYC, and GPX4 in DHL10 (left) and OCI-Ly1 (right) cells treated with (B) erastin (600 nM) + zotatifin (12.5 nM) or (C) RSL3 (50 nM) + zotatifin (12.5 nM) for 24 hours. Data from 2 independent experiments. D) Oxidized BODIPY C11 fluorescence measured by flow cytometry in SU-DHL10 (left) and OCI-Ly1 (right) cells treated with zotatifin alone or with ferroptosis inducers (erastin, RSL3, DMF) or inhibitor (NAC). Statistical analysis used one-way ANOVA with Bonferroni multiple comparison test (*P < 0.05, **P < 0.01, ***P < 0.001, ****P < 0.0001). E-F) ROS measurement by flow cytometry in SU-DHL10 (left) and OCI-Ly1 (right) cells stained with (E) MitoSox and (F) DHE, treated as above. Statistical analysis used one-way ANOVA with Bonferroni multiple comparison test (*P < 0.05, **P < 0.01, ***P < 0.001, ****P < 0.0001).

### NRF2 is a direct translational target of rocaglates and mediates sensitivity to ferroptosis

Genome wide CRISPR/Cas9 screen associated NRF2 activation with Rocaglate resistance (4) but its protective mechanisms in this context are not well defined. NRF2 activation drives protection from ferroptosis (33), and we therefore hypothesized translational NRF2 loss could explain both the published CRISPR screen and our results with ferroptosis inducers. NRF2 was reported to decline in response to rocaglates in metastatic sarcoma models, enabling the activity of ROS-inducing compounds (34). Analysis of CCLE cell lines revealed significant resistance to CR-1-31B with increasing expression of *NFE2L2*, encoding NRF2, or *HMOX1*, encoding the NRF2-regulated protein Heme Oxygenase-1 (HO-1). In contrast, *KEAP1*, the encoding Kelch-like ECH-associated protein 1 (KEAP1) that promotes NRF2 degradation exhibited the opposite trend (**Fig. 4SA**). To further investigate rocaglate impact on NRF2 in DLBCL, we assessed NRF2 levels in response to zotatifin, revealing decline similar to MYC (**Fig. 4A**). HO-1 declined in parallel. To confirm this is due to eIF4A1 inhibition, we utilized CRISPR-engineered HAP1 cells expressing eIF4A1-F163L, which eliminates zotatifin binding to eIF4A1 without impacting its function in translation (8,18,35). While zotatifin downregulated NRF2 protein in wild-type and Cas9 control (F163F) HAP1 cells, the effect was abolished in eIF4A1-F163L mutants (**Fig. 4B**). DDX3, which is unmodified in these cells, is a secondary RNA helicase target of rocaglates, and its inhibition could explain the observed NRF2 increase in F163L cells (36). NRF2 loss in response to zotatifin is therefore eIF4A1-dependent.

**Figure 4:**
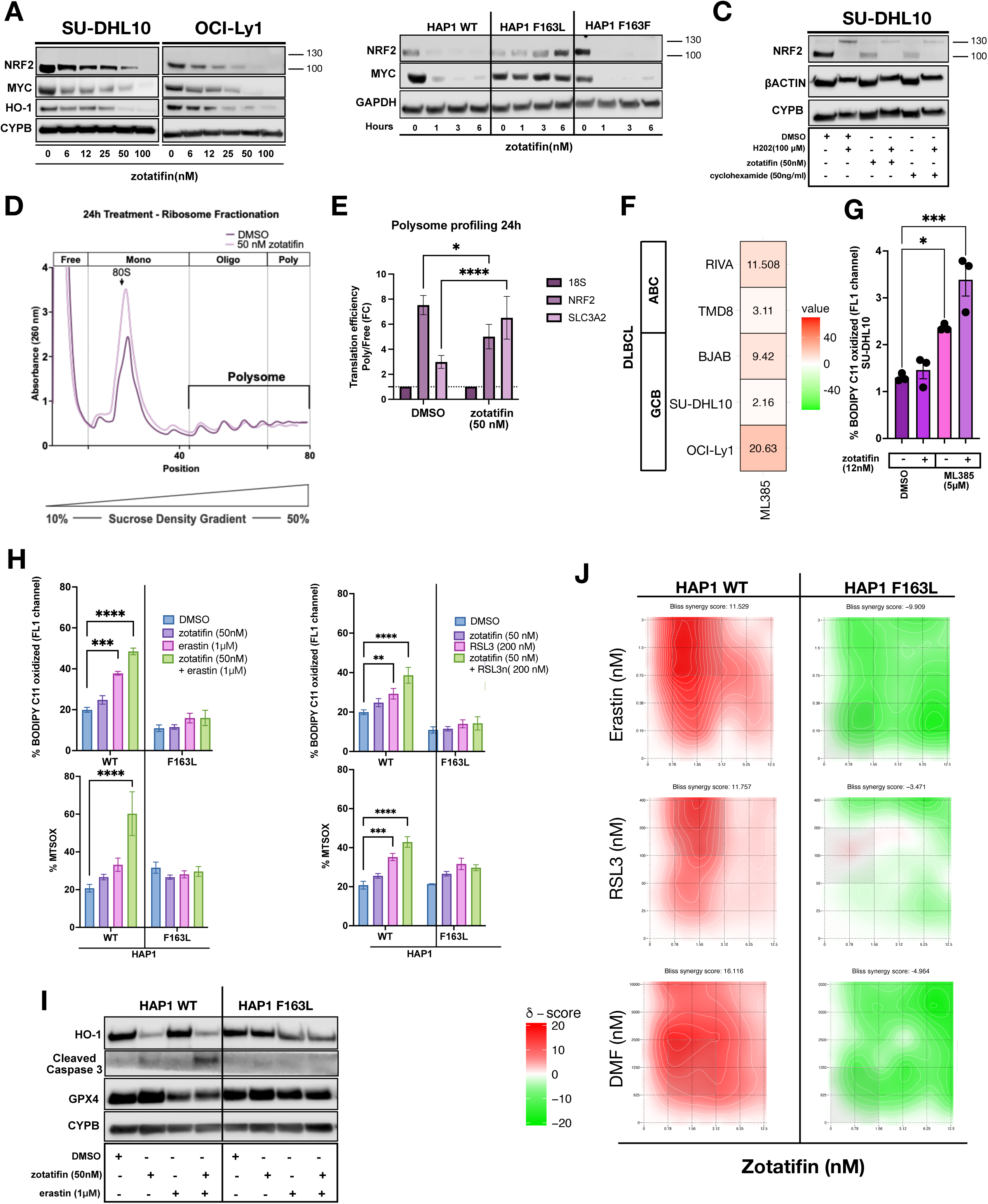
A) Immunoblot analysis of SU-DHL10 (left) and OCI-Ly1 (right) cells treated with zotatifin at concentrations of 0, 6.25, 12.5, 25, 50, and 100 nM for 24 hours for NRF2, MYC, and HO-1. Data are representative of 2 independent experiments. B) Immunoblotting of HAP1 cells (WT, F163L, F163F) for NRF2 and MYC treated with 100 nM zotatifin for various durations (0, 1, 3 6 hours). Data are representative of 2 independent experiments. C) Immunoblot of SU-DHL10 treated with H_2_0_2_ (100 μM) for 2 hours alone or in combination with 100 nM zotatifin or 50 ng/mL cycloheximide for NRF2. Data are representative of 2 independent experiments. D) Polysome profiling of SU-DHL10 cells treated with vehicle or 50 nM zotatifin for 24 hours. Polysome data are mean ± SEM of 2 independent experiments. E) mRNA translation efficiency in SU-DHL10 cells treated with 50 nM zotatifin for 24 hours, measured as the ratio of polysome-associated mRNA to ribosome-free and monosome-associated mRNA. Data represent mean ± SEM of 2 independent experiments. Unpaired t tests were performed to determine statistical significance (^∗^p ≤ 0.05, ^∗∗∗∗^p ≤ 0.0001). F) Bliss δ synergy score heatmap for zotatifin in combination with ML385 in RIVA, TMD8, BJAB, SU-DHL10, and OCI-Ly1 DLBCL cell lines. A Bliss δ synergy score > 10 indicates synergy, between −10 and 10 suggests an additive effect, and < −10 indicates antagonism. G) Oxidized BODIPY C11 fluorescence measured by flow cytometry in SU-DHL10 cells treated with zotatifin and ML385. Statistical analysis: one-way ANOVA with Bonferroni multiple comparison test. Data are mean ± SEM of three independent experiments. *P < 0.05, ****P < 0.0001 vs. DMSO. H) Lipid peroxidation (top) and ROS measurement (bottom) with BODIPY C11 and MitoSOX in HAP1 cells WT(left) F163L (right) treated with zotatifin and erastin (left) or RSL3 (right). Data represent mean ± SEM of 3 independent experiments. Statistical analysis was performed using repeated measures two-way ANOVA with Sidak’s multiple comparison test (**P < 0.01, ***P < 0.001, ****P < 0.0001). I) Immunoblot of HAP1 cells (WT and F163L) treated with 1 µM erastin and 50 nM zotatifin for 24 hours for HO-1, cleaved caspase 3, and GPX4. Data are representative of 2 independent experiments. J) Bliss synergy plots for zotatifin combined Erastin, (top) RSL3 (center) and DMF (bottom) with zotatifin in HAP1 WT (left) and F163L(right) and A549 (right) and H460 (left). A Bliss δ synergy score > 10 indicates synergy, between −10 and 10 suggests an additive effect, and < −10 indicates antagonism.

For further investigation, we compared rocaglate treatment to cycloheximide, which specifically targets NRF2 translation rather than transcription (37). Unlike Actinomycin D, cycloheximide (CHX) can reverse the NRF2 response to H_2_O_2_ (37). Mikac et al. identified two distinct molecular weights of NRF2 (105 and 130 kDa) in lung cancer cell lines (38). The 105 kDa form of NRF2 is stable and resistant to Keap1-Cul3-mediated degradation, potentially explaining the factor’s constitutive activity in these cells. D1Z9C antibodies recognize the 130 kDa form, which represents the Keap-dependent transient NRF2 form induced by oxidative stressors. We found both zotatifin and CHX impaired NRF2 expression of both the 100 and 130 kDa forms, reversing H_2_O_2_ effects (**Fig. 4C**). We next assessed translational cell fractions by polysome profiling after treatment with vehicle or zotatifin (50 nM x24 hours, **Fig. 4A**). These studies revealed zotatifin reduced translation efficiency of NRF2 (**Fig. 4D-E**) while increasing efficiency for the up-regulated target SLC3A2. We also tested the direct NRF2 inhibitor ML385 (39) and found synergy or additivity with zotatifin across DLBCL cell lines, and the combination significantly increased lipid peroxidation compared to either alone (**Fig. 4F-G**). In HAP1 cells, lipid peroxidation, ROS, synergy, and decline of GPX4 and the NRF2 target HO-1 were abrogated in eIF4A1-F163L mutant cells compared to WT for erastin, RSL3, and DMF (**Fig. 4H-J**). Collectively, these results are consistent with NRF2 translational loss as the basis for rocaglates’ sensitization of target cells to inducers of ferroptosis.

### In vivo efficacy of zotatifin combined with the ferroptosis inducer imidazole ketone erastin

To evaluate synergy in vivo, we used the in vivo optimized analog imidazole ketone erastin (IKE, **Fig. 5A**) (40). We employed a patient-derived xenograft (PDX) model of GCB-DLBCL (ProXe DFBL-75549-R2) (41). Initially, 10 mice per group were treated with IKE at 40 mg/kg intraperitoneally (IP) 5 times per week (40), zotatifin at 1 mg/kg IP twice weekly (18), or the combination. However, by day 10, four mice in the combination group experienced significant weight loss meeting ethical endpoints and had to be sacrificed (**Fig. 5B**). Gross examination suggested liver toxicity indicated by pale discoloration of the organ. We therefore adjusted dosing in the combination group to twice weekly for IKE and once weekly for zotatifin, and the experiment continued without further host toxicity. Single-agent IKE had no significant impact on tumor volume or overall survival compared to vehicle, while zotatifin alone prolonged survival but did not achieve significant tumor volume (TV) reduction during the period when vehicle-treated mice were still alive (**Fig. 5C-D**). In contrast, combination therapy dramatically reduced tumor volume and significantly prolonged overall survival, even with four animals lost early due to toxicity. After dose adjustment, mice in the combination group experienced no further toxicity and had minimal tumor burden at the end of the experiment (day 40, **Fig. 5D**). Rocaglate synergy with ferroptosis inducers is therefore exploitable in vivo but required dose modification compared to the single agents to prevent host toxicity.

**Figure 5:**
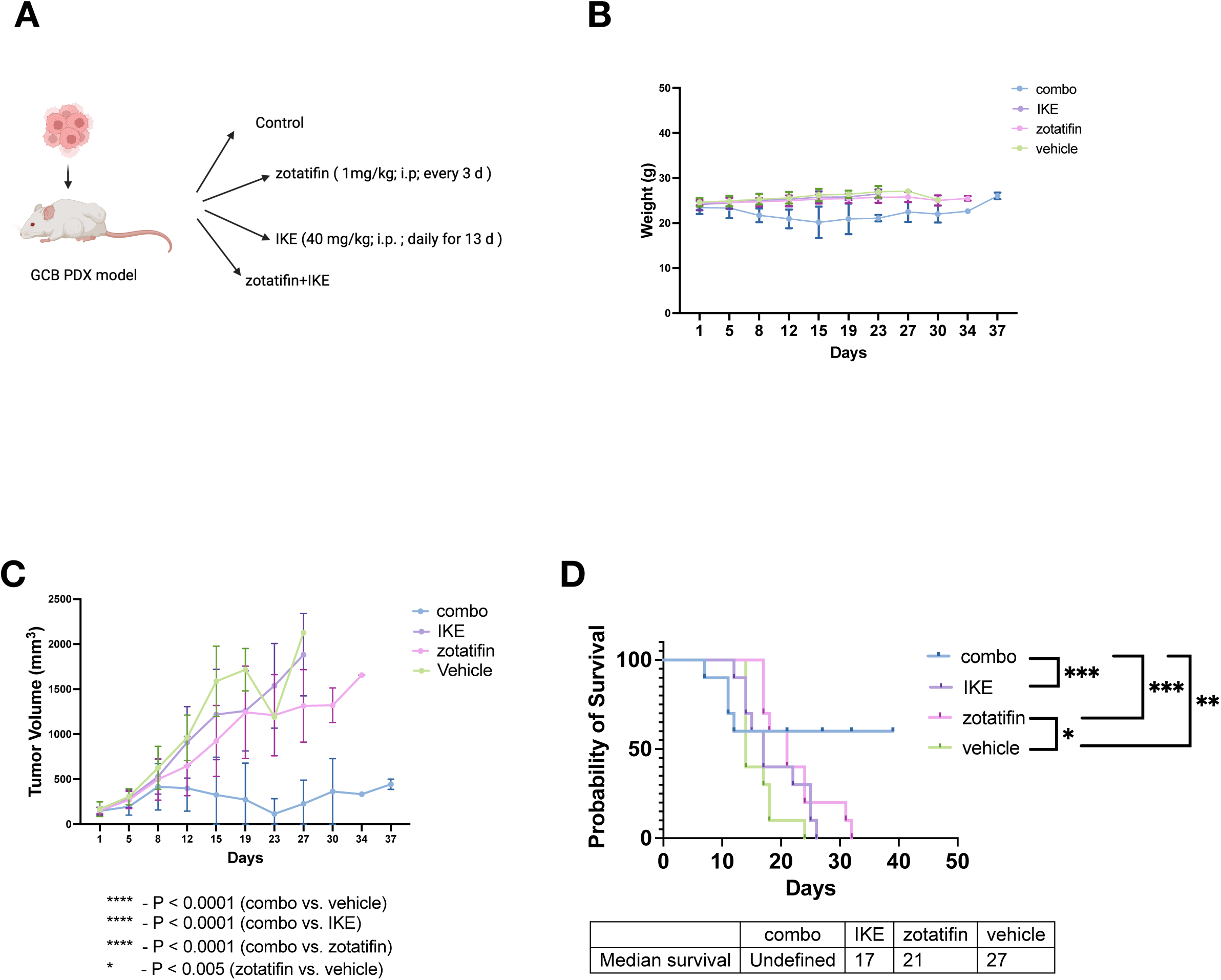
A-C) Tumor growth and survival in a DLBCL GCB PDX mouse model. A) 40 DLBCL GCB PDX tumors were implanted in NOD scid gamma mice, randomized into four groups of 10 mice each: control, zotatifin (1 mg/kg, IP injection twice a week), IKE (40 mg/kg, IP injection five days a week for 13 days), and combination (zotatifin + IKE). When average tumor volume (TV) exceeded 150 mm³, zotatifin dosing was reduced to twice a week, and IKE dosing to once a week. B) Average weights ± SEM measured twice weekly. C) Average tumor volume ± SEM measured twice weekly via ultrasound. Data represent mean ± SEM of five independent experiments. Statistical analysis was performed using repeated measures two-way ANOVA with Sidak’s multiple comparison test (*P < 0.05, **P < 0.01, ***P < 0.001, ****P < 0.0001). D) Kaplan-Meier curves showing survival probability for all treatment groups for five independent experiment. ***P < 0.001, **P < 0.01, *P < 0.05 (Mantel-Cox).

### Zotatifin primes large B-cell lymphomas for killing by CD19 CAR-T cells

Prior studies have shown CD8+ T cells activated by immunotherapy cause ferroptosis in tumor cells (16). Release of interferon gamma (IFNγ) downregulates the system X_c_^-^ components SLC3A2 and SLC7A11 in targets, hindering cystine uptake and protection from lipid peroxidation (16,42). We therefore tested whether rocaglate synergy with ferroptosis inducers would extend to IFNγ, revealing strong zotatifin-IFNγ synergy in DLBCL cell lines (**Fig. 6A**). IFNγ is one of the main pro-inflammatory cytokines secreted by CD8+ effectors in CAR-19 products and cross activated by them in lymphoma microenvironments to drive clinical responses (43,44). We therefore reasoned CAR-19 therapy might synergize with rocaglates in inducing ferroptosis and tumor eradication. We first needed to determine whether zotatifin can be directly combined with CAR-T cells therapeutically without toxicity to the immune cells themselves. Employing second-generation CD19-directed CAR T cells with a CD28 costimulatory domain, we found zotatifin significantly reduced viability, including complete death at higher concentrations and exposure times, making co-treatment therapeutically incompatible (**Fig.6B**). We therefore pretreated SU-DHL10 and OCI-Ly1 target cells with zotatifin (12.5nM) for 24 hours, followed by drug washout before co-culturing with varying concentrations of CAR-T cells (**Fig.6C**). Zotatifin-pretreated cells were significantly more responsive to CD19-CD28-z CAR T cells than DMSO-pretreated cells in both, and we assessed also A20 murine lymphomas treated with strain-matched CD19-CD28-z CAR T cells (**Fig. 6D**) (45). We further confirmed this heightened sensitivity using a human 4-1BB-z CAR-T construct (**Fig. S5A**). Sensitization to cytotoxicity by zotatifin pretreatment was most pronounced at lower effector-target (E:T) ratios. Consequently, we assessed the levels of ROS since lipid peroxidation staining presented challenges due to overlapping flow cytometry spectra. We found elevated levels of reactive oxygen species (ROS) at lower E:T ratios of CARs (0.3:1-1.25:1), in cells pre-treated with zotatifin (**Fig. S5B**).

**Figure 6.**
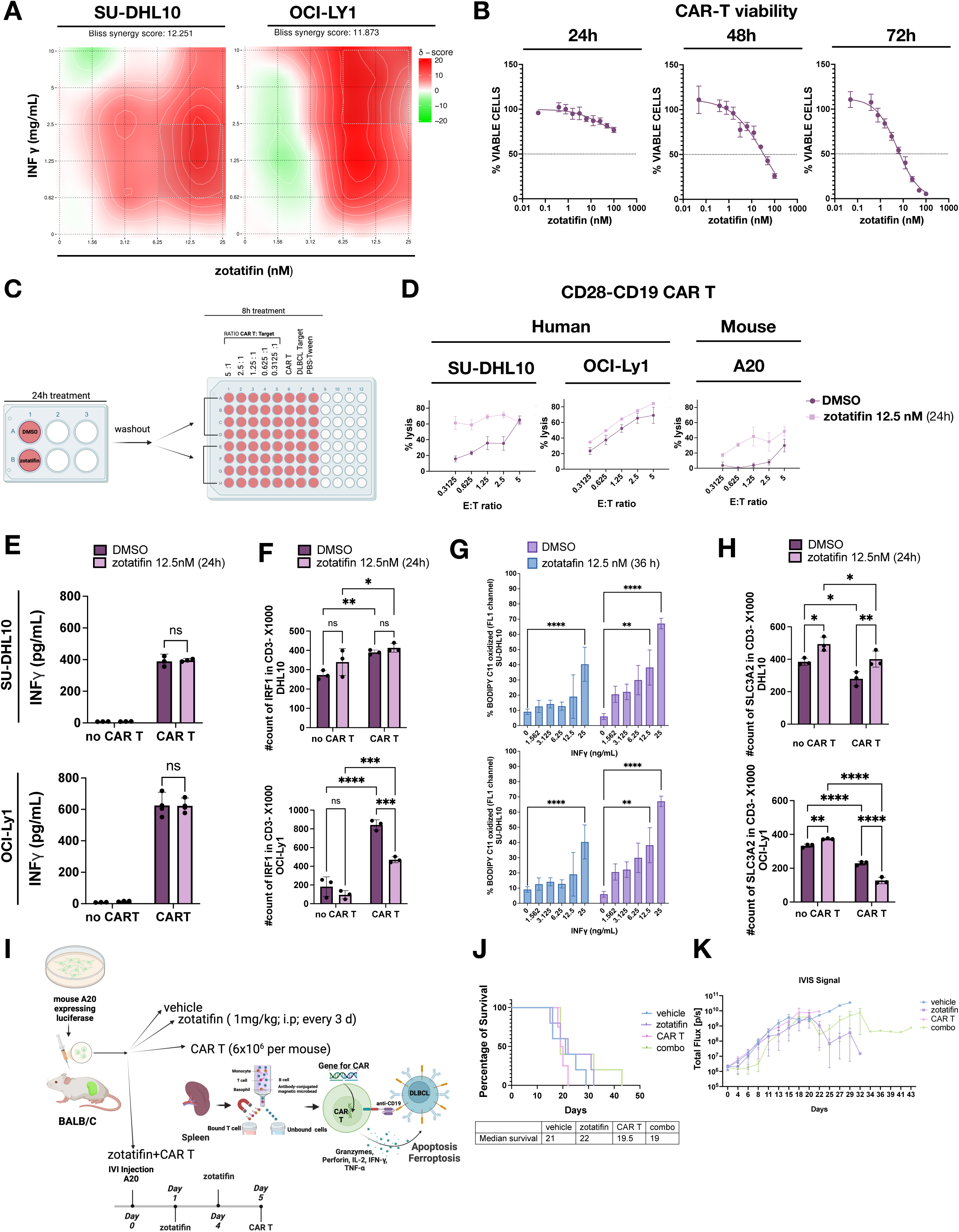
(CAR T): A) Bliss synergy plots for zotatifin combined with IFN γ in SU-DHL10 (left) and OCI-Ly1 (right) cells. A Bliss δ synergy score > 10 indicates synergy, between −10 and 10 suggests an additive effect, and < −10 indicates antagonism. B) ATP levels in CD28-CD19 CAR T AD114-derived cells at 24h(left), 48h (center), and 72h (right). C) Schematic of co-culture experiment. D) 8h co-culture of RD114-derived CD28-CD19 CAR T cells with SU-DHL10 (left) and OCI-Ly1 (center) or mouse CD28-CD19 CARs with A20 Luciferase (right), CFSE+ pretreated DLBCL (DMSO or 12.5 nM zotatifin). Measured by flow cytometry (see schematic C). E) IFN γ levels from co-culture (see schematic C) with SU-DHL10 (top) and OCI-LY1 (bottom) cells at a 1.25 (CARs):1 (DLBCL) ratio. Data represent mean ± SEM of three experiments. Statistical analysis was performed using repeated measures two-way ANOVA with Sidak’s multiple comparison test. F) 8h co-culture of CD28-CD19 CAR T cells (see schematic C) with SU-DHL10 (top) and OCI-Ly1 (bottom) cells, fixed, permeabilized, and probed with CD3 and IRF1. Measured by flow cytometry at a 1.25 (CARs):1 (DLBCL) ratio. Data represent mean ± SEM of three experiments. Statistical analysis was performed using repeated measures two-way ANOVA with Sidak’s multiple comparison test *P < 0.05, **P < 0.01, ***P < 0.001, ****P < 0.0001). G) Oxidized BODIPY C11 fluorescence in SU-DHL10 (top) and OCI-Ly1 (bottom) cells pretreated with DMSO or 12.5 nM zotatifin for 36h, then treated with IFN γ (0-25 ng/mL) for the last 8h. Data represent mean ± SEM of three experiments. Statistical analysis was performed using repeated measures two-way ANOVA with Sidak’s multiple comparison test **P < 0.01, ****P < 0.0001). H) 8h co-culture of axi-cel (see schematic C) with SU-DHL10 (top) and OCI-Ly1 (bottom) cells, fixed, permeabilized, and probed with CD3 and SLC3A2. Measured by flow cytometry at a 1.25 (CARs):1 (DLBCL) ratio. Data represent mean ± SEM of three experiments. Statistical analysis was performed using repeated measures two-way ANOVA with Sidak’s multiple comparison test *P < 0.05, **P < 0.01, ***P < 0.001, ****P < 0.0001). I) Schematic of in vivo CAR T experiment. 40 BALB/c mice were engrafted with A20-expressing luciferase cells (1×10^6^ cells/mouse) via tail injection and randomized into four groups of 10 mice each: vehicle, zotatifin (1 mg/kg, IP every three days), CAR T, and combo. CAR T cells (6×10^6^ cells/mouse) were injected at day 5 into CAR T and combo groups. CAR T cells were collected from spleen 14 days before injection. J) Kaplan-Meier curves showing survival probability for all treatment groups from five independent experiments. K) Average total flux signal (p/s) over time for mice implanted with luciferase-expressing A20 cells. Data represent mean ± SEM of five independent experiments. Statistical analysis was performed using repeated measures two-way ANOVA with Sidak’s multiple comparison test.

To evaluate IFNγ release upon ferroptosis induction mediated by CAR T, given the potential for zotatifin to hinder CAR T function, we assessed its levels in the co-culture (**Fig. 6E**) at E:T ratios of 1.2:1. We found CARs triggered IFNγ accumulation (**Fig. 6E**) as well as IRF1 induction (**Fig. 6F**) in targets, with no notable difference between DMSO and zotatifin pretreatment. Therefore, neither production of IFNγ nor activation of STAT1-mediated downstream signaling were significantly impacted from treating lymphomas with zotatifin before CAR-T cell challenge. Interestingly, recent findings (18) suggest zotatifin itself may prompt a cell-intrinsic IFNγ response. We therefore assessed IFNγ production by target DHL-10 cells during zotatifin exposure (**Fig.S5C**). Though IFNγ secretion occurred in response to the compound, the level of production was 10-20-fold lower than what occurred during co-culture with CAR-T cells (compare Y-axis scale **Fig. 6E to S5C**). In addition, washout of drug for 8 hours completely eliminated increased IFNγ, and therefore all increase in levels of the cytokine curing co-culture likely came from the CAR-T effectors. Finally, our findings suggest that ferroptosis induction during CAR-T co-culture is likely due to IFNγ, as treatment with IFNγ for 24 hours, with or without zotatifin washout, indicated IFNγ-induced ferroptosis, with a particularly notable synergy between IFNγ and zotatifin in inducing ferroptosis (**Fig. 6G**). Overall, these results implicate induction of inherent sensitization to the effects of the CAR-T cells by the zotatifin pretreatment, consistent with impaired response to ferroptotic stress during engagement by the immune effectors.

Zotatifin induces protein expression of SLC3A2 (5) while IFNγ secreted by CD8+ cells is known to promote its transcriptional repression (16). We therefore conducted the same coculture experiment at E:T ratios of 1.2:1 and assessed SLC3A2 protein by flow cytometry on the target (CD3-) cells. We observed that while zotatifin alone stimulates SLC3A2 expression that persists even after drug washout, the effect was noticeably attenuated during CAR-T coculture (**Fig. 6H**). Therefore, as we observed with pharmacologic inducers of ferroptosis, the protective response against ferroptosis induced by zotatifin is insufficient to overcome a critical loss of NRF2, sensitizing target cells to CAR-T-mediated cytotoxicity.

Finally, to determine the in vivo sensitivity of zotatifin combined with CAR T cells, we engrafted A20 cells (BALB/c lymphoma cells derived from spontaneous reticulum cell neoplasm) expressing luciferase IV via tail injection (1×10^6^cells per mouse) in syngeneic BALB/c mice. To mitigate the impact of zotatifin on CAR T cells while leveraging zotatifin’s enhanced susceptibility to ferroptosis, considering the half-life of zotatifin in mice is 10.9 hours (8), treatment began on day 5 after physiological washout of zotatifin (**Fig. 6I**). We injected 6×10^6^ CAR T cells per mouse. Tumor volume at D15 (10 days post CAR infusion) was transiently improved with the combination, and mice in the combo group achieved increased overall survival surviving for more than 40 days (**Fig. 6J-K, S5D**). Particularly on day 8, following the early injection of CAR T cells, we found that the combination significantly decreased tumor progression (**Fig. S5E**). These results suggest optimization of experimental conditions might better capture improved CAR-T efficacy after zotatifin pre-treatment in vivo. However, due to questions that arose regarding zotatifin’s availability for short-term clinical translation related to wind-down of its developer, eFFECTOR Therapeutics, we were unable to justify expense and animal usage for further such experiments (see discussion).

## Discussion

Preclinical studies of cap-dependent translation inhibitors highlight therapeutic potential across multiple tumor types, rooted in lost expression of key oncoproteins, many of which are otherwise undruggable. Though varying in mode-of-action, essentially all these compounds bind to eIF4A1, the ATP-dependent enzymatic core of the cap-binding/mRNA unwinding translation initiation complex eIF4F. Rocaglates, including both natural and synthetic compounds, have been the most extensively studied, revealing sequence-dependent clamping of the target to RNA, a highly unusual molecular pharmacology (46). Despite this growing body of research dating back to the 2000s, only one eIF4A1 inhibitor, zotatifin, has reached clinical evaluation. Aggressive B-cell lymphomas have arguably the strongest body of published support for therapeutic potential of these compounds (9–11), but due to larger markets, zotatifin’s developer eFFECTOR Therapeutics focused clinical trials in solid tumors, including KRAS-mutant NSCLC and hormone receptor-positive breast cancers. Data have not been published, but eFFECTOR’s recently announced corporate wind-down indicates disappointing results (47). Fortunately, a variety of rocaglates and pateamine derivatives remain candidates for clinical development to permit testing in, we argue, the much more therapeutically fertile ground of B lymphocyte-derived malignancies. Here, we exploited zotatifin’s favorable pharmacologic properties to further investigate the therapeutic potential of eIF4A1 inhibition in lymphoma. Combined RNA-seq and mass spectrometry revealed decoupling between transcriptional and translational responses. A recent study in metastatic osteosarcomas observed comparable stress responses in transcriptomes and translatomes in response to CR-1-31B, but did not specifically compare the up- or downregulation of individual genes in the proteome and translatome (34). While we have previously noted this effect (5), our study is the first to demonstrate it on a broader scale, with zotatifin limiting translation of upregulated mRNAs. Mechanistically, we highlight NRF2, a central regulator of the antioxidant responses, as a direct translational target of rocaglates. Yabaji et al. reported that rocaglates up-regulated NRF2 transcriptional targets in macrophages during Mycobacterium tuberculosis infection (48) but did not assess changes in NRF2 protein levels, and the infection context may influence these results differently. In osteosarcomas, targeting eIF4A1 hindered NRF2 expression and reduced metastases, and the oxidative stress inducer tert-butyl hydroquinone (tBHQ), when combined with CR-1-31B, synergistically augmented cell death (34). Our results refine the understanding of NRF2 loss as a direct translational rocaglate consequence and expand its implications to ferroptosis sensitization and additional tumor types. Previous CRISPR/Cas9 screens linked NRF2 activation to rocaglate resistance (4) but mechanisms were unclear. Our findings findings underscore the importance of NRF2 loss in enhancing sensitivity to ferroptosis and suggest that targeting its translation with eIF4A1 inhibitors could improve therapeutic strategies in cancers with high NRF2 activity.

In vivo experiments demonstrated the synergistic potential of combining zotatifin with ferroptosis inducers for DLBCL treatment. Using IKE in a patient-derived xenograft model, we found that single-agent IKE had no significant impact on tumor volume or survival, and zotatifin alone extended survival without notable tumor reduction. The combination, however, led to significant improvements in both outcomes. However, initial dosing resulted in significant toxicity, which previously was seen also for single-agent IKE at higher doses (40), necessitating adjustments to complete our study. Our finding that zotatifin primes DLBCL cells for enhanced killing by CAR-T cells is an exciting potential avenue to leverage CAR-19 efficacy but requiring further optimization prior to clinical translation. Previous studies have shown that CD8+ T cells activated by immunotherapy induce ferroptosis in tumor cells via IFNγ release, which downregulates critical cystine transporters (16). Recent research has also highlighted the synergy between rocaglates and IFNγ in triple-negative breast cancer (18). Building on these findings, our results demonstrated a strong synergy between zotatifin and IFNγ in lymphoma cell lines, suggesting zotatifin can sensitize tumors to CAR-T cell-mediated ferroptosis in a manner similar to pharmacologic inducers. Importantly, however – and not surprisingly – zotatifin was directly toxic to CAR-T products. In a co-culture, setting, we were able to establish sensitization to CAR-19 killing through zotatifin pre-treatment, with CAR-T cells applied to targets rapidly after drug washout. These results occurred across multiple systems and CAR constructs, especially at lower effector-to-target ratios. In vivo, drug half-lives made washouts and timing more challenging, and we achieved only a transient sensitization, although encouragingly co-treated animals achieved prolonged survival. The favored approach in our opinion would be to engineer effector populations with the F163L rocaglate-resistance mutation through CRISPR modification concomitantly with CAR introduction. This intriguing but technically challenging cell-engineering effort was outside our current scope.

In summary, this study uncovers a novel mechanism by which zotatifin enhances ferroptosis sensitivity and augments CAR-T cell therapy efficacy in DLBCL. By elaborating the role of NRF2 loss and the decoupling of transcriptional and translational responses, we highlight zotatifin’s unique ability to prime tumor cells for ferroptotic stress, ultimately increasing their susceptibility to CAR-T cytotoxicity. Results suggest combining eIF4A1 inhibitors with ferroptosis inducers could be a promising strategy to overcome therapeutic resistance in relapsed or refractory DLBCL. Future research should focus on optimizing the dosing and timing of these combinations to maximize their clinical potential, particularly in cancers with high NRF2 activity.

## Methods

### Reagent

All reagents used in the experiments are listed in **Supplementary Table S1**.

### Cell lines

All cell lines were routinely verified by STR fingerprinting and confirmed mycoplasma negative using the PlasmoTest Kit (Invivogen: REP-PT1). SU-DHL-10, RIVA, BJAB, TMD8 were grown in RPMI (Fisher Scientific, 11-875-119) with 10% FBS (VWR) and penicillin/streptomycin (P/S, VWR) (R10 complete media). OCI-Ly1 was grown in IMDM (Fisher Scientific, 12440053) with 20% FBS and P/S. Cell lines were purchased by American Type Culture Collection (ATCC), Manassas, VA, USA, RIVA was purchased from DSMZ. Cells were maintained at 37 °C in a 5% CO_2_ humidified incubator.

### Transcriptomic Analysis in the CCLE Dataset

Transcriptomic data and drug sensitivity (EC50) values were from the Cancer Cell Line Encyclopedia (CCLE) (https://depmap.org/portal/ccle)(26), including sensitivity to the synthetic rocaglate CR-1-31B (**Table 1**). Linear regression analysis was employed to model the relationship between gene expression and EC_50_, with slope indicating the change in drug sensitivity per unit increase in gene expression. The statistical significance of the regression model was assessed using p-values. All analyses were performed using DepMap Data Explorer and visualized with RStudio (R 4.4.1, Posit, https://posit.co/download/rstudio-desktop/).”

### Engineered Cells

To achieve overexpression of SLC3A2, we utilized TET-inducible pLV [Exp]-Puro TRE>hSLC3A2[NM_001012661.1] (VB211209-1178dwk, Vector Builder), along with the helper vector pLV[Exp]-CMV>tTS/rtTA/Hygro (VB010000-9369xhm, Vector Builder).

### CAR-T cell production

#### Human

Human CAR-19 cells with the 4-1BB costimulatory domain were purchased from BPS Bioscience (Cat #: 78171-1) and cultured per manufacturer’s instructions. Human CAR-19 cells with the CD28 costimulatory domain were produced by infecting PBMCs with retroviral supernatants from the SFG-h1928z RD114 producer line, as described previously (45). Briefly, stable RD114 producer cells were generated by infection of retroviral supernatants from SFG-h1928z transfected H29 cells. Supernatant from RD114 producers was collected and filtered two days after seeding 5×10^6^ cells in a 15 cm dish. Takara Retro-X concentrator was then used to concentrate filtered SFG-h1928z virus following the manufacturer’s instructions and resuspended in 1/50^th^ volume of harvested supernatant in PBS. Transduction and subsequent culture of donor PBMCs with SFG-h1928z virus was carried out as previously described (49).

#### Mouse

All mouse studies were performed under the approval of the University of Miami Institutional Animal Care and Use committee. The mice were housed in pathogen free conditions. All cages are handled under animal transfer stations (Biological Safety Cabinets). Mice are housed in individually ventilated, double sided, racks and are changed weekly. Mouse CAR-19 cells were generated first by isolating T cells from spleens of BALB/c mice followed by CD3 negative selection (Stemcell Technologies #19851A). T cells were activated and transduced with m1928z-mCherry vector as previously described (45).

#### CAR T co-culture

Cells were pretreated with DMSO and 12.5nM of zotatifin for 24h. After 24h cells were washed and coculture were prepared in 96well for coculture with CARs as follows: 1×10 ^5^ DLBCL cells (SU-DHL10 or OCI-Ly1) were stained with CFSE (**Table S4**) in 1mL of naked media for 15 minutes at 37°C. Cells were than washed, centrifuge and resuspended in complete media for 10 minutes at 37°C. DLBCL cells were than resuspended in 3mL of complete media. 5 ×10^5^ CARs cell were separated from the beads with a magnetic column centrifuge and resuspended in 1mL of complete media. 100μL CARs cell were serially diluted in complete media from wells 1-6. 100μL CFSE+ cells were distributed from column 1-5 to obtain obtain ratio (5:1, 2.5:1, 1.25:1,0.63:1, 0.31:1) with CARs cell. 100μL cells were add to column 7 as a negative control. Column 8 were filled with 100μL DLBCL cells and 100μL PBS-0.02% Tween as a positive control for lysis. Co-colture experiments were performed for 8h.

#### Flow Cytometry for CAR T staining

Co-culture wells (96-well plates) were first washed with autoMACS rinsing buffer containing BSA (Miltenyi Biotec, #130-091-222 and #130-091-376). Cells were then treated with FcR-blocking reagent (Miltenyi Biotec, #130-092-575) and stained with fluorescently conjugated antibodies, including Invitrogen Antibody listed in **Supplementary Table S4**. After staining, cells were fixed with 1% formaldehyde solution (Thermo Fisher Scientific, #047392-9M). Flow cytometry data acquisition was performed using a Cytek Aurora, and the data were analyzed with FlowJo v.10 software, applying default parameters (iterations = 1000, perplexity = 30, θ = 0.5).

#### Enzyme-Linked ImmunoSorbent Assay (ELISA) and multiplex cytokine array

Condition media or cellular protein lysates were quantified, and equal amount of protein loaded per well of a standard 96-well plate, including three technical replicates per biological replicate. ELISA kits specific for mouse TNF-alpha and IFN-γ (R&D Systems, # DTA00D, and # DIF50C) were used according to manufacturer protocol to quantify cytokines levels.

#### TMT-pSILAC (tandem mass tag-pulse stable isotope labeling with amino acids in cell culture mass spectrometry)

Each cell line was tested in three independent replicates per condition: zotatifin (12.5 nM for 24 hours) or DMSO. For pSILAC experiments, cells were grown in light (R0K0) SILAC media for 7 days and then pulsed with heavy (R10K8) SILAC media (Thermo Fisher Scientific) for the final 8 hours of zotatifin treatment. Cells were lysed using polysome lysis buffer (0.3 M NaCl, 15 mM MgCl2·6H2O, 15 mM Tris-HCl pH 7.4, 1% Triton X-100, 0.1 mg/ml cycloheximide, 100 units/ml RNase inhibitor).

Heavy labeling for mass spectrometry was performed by Bioinformatics Solutions Inc. (Waterloo, Ontario, Canada), as previously described. Samples were reduced, alkylated, digested, and labeled using the TMT10plex™ Isobaric Label Reagent Set following the manufacturer’s instructions. High-pH reversed-phase high-performance liquid chromatography (HPLC) fractionation was carried out using a Waters XBridge C18 column. The resulting fractions were prepared for MS analysis on a Thermo Scientific Orbitrap Exploris™ 240. Mass spectrometry data were processed using PEAKS X+ (v10.5, Bioinformatics Solutions Inc.). Detailed methods can be found in our previous publication (5).

#### Ribosome density profiling

Ribosome density profiling was conducted as follows: Cells were treated with 0.1 mg/ml of cycloheximide for the last 10 minutes of treatment, followed by ice-cold washes with PBS containing cycloheximide (0.1 mg/ml). Subsequently, cells were lysed in polysome lysis buffer consisting of 0.3 M NaCl, 15 mM MgCl2.6H2O, 15 mM Tris-HCl pH 7.4, 1% Triton X-100, 0.1 mg/ml cycloheximide, and 100 units/ml RNase inhibitor. After centrifugation (twice at 10,000 g for 5 minutes at 4 °C) to remove cellular debris, samples were loaded based on equal total RNA onto a 10%–50% sucrose gradient and subjected to ultracentrifugation (187,813 g for 1.5 hours at 4 °C) using a SW 41 Ti rotor (Beckman Coulter). Subsequently, samples were fractionated into 1 ml fractions and collected using the BR-188 density gradient fractionation system (Brandel). Total RNA was isolated from each fraction through phenol-chloroform extraction and ethanol precipitation following proteinase K treatment.

#### Immunoblot

SDS-PAGE was conducted utilizing BoltTM 4%–12% Bis-Tris Plus pre-made gels from ThermoFisher Scientific, employing the Mini Gel Tank system also from ThermoFisher Scientific. Subsequently, proteins were transferred onto 0.2 μm Immuno-Blot® PVDF membranes (Bio-Rad) using the BoltTM Mini Blot Module from ThermoFisher Scientific, following the manufacturer’s protocols. Chemiluminescent signals were captured using a digital chemiluminescent imaging system. Densitometry analysis was carried out on non-saturated signals utilizing ImageJ (NIH), and normalization was performed with respect to the loading control. All antibodies are listed in **Supplementary Table S2.**

#### qRT-PCR

First-strand cDNA synthesis was performed using the Maxima H Minus cDNA Synthesis Master Mix (ThermoFisher Scientific, #M1661), according to the manufacturer’s protocols. qRT-PCR was performed using a StepOnePlus^TM^ Real-Time PCR System (ThermoFisher Scientific). Relative changes in expression were calculated using the comparative Ct (ΔΔCt) method. 18S rRNA was measured as an internal control for changes in RNA levels. All qPCR reactions were performed using PowerUp^TM^ SYBR® Green Master Mix (ThermoFisher Scientific, #A25742) unless otherwise stated. Primers are listed in **Supplementary Table S3**

#### Synergy assay

For synergy investigation, we employed different DLBCL cell lines (OCI-Ly1, SU-DHL10, RIVA, TMD8, BJAB) and test synergy using the web application SynergyFinder and applying Bliss model (50), where values >10 indicate synergism, from −10 to 10 – additivity, values <-10 – antagonism.

#### Cell viability and apoptosis assay

Cell viability was assessed by luminescence measurements using CellTiter-Glo® and RealTime-Glo MT viability assays (Promega). Annexin V-fluorescein isothiocyanate/propidium iodide (Annexin V-FITC/PI, BD Pharmingen™) double staining was used to quantify apoptosis in SU-DHL10 overexpressing CD98hc. Cells were incubated with 0, 12.5nM of zotatifin for 24h and 48h hours in doxycycline-induced (800ng/mL) and not induced, then collected. Cells were washed twice with ice-cold PBS and resuspended in 1× Binding Buffer at a concentration of 1×10^6^ cells/ml. 5 μl of Annexin V-FITC and 5 μl of propidium iodide were added to 100 μl of the cell suspension. After 15 minutes of incubation, 400 μl of Binding Buffer were added to each cell suspension. Cells were analyzed by flow cytometry within 1 hour.

#### Glutathione assay

The GSH/GSSG assay was conducted using the Promega GSH/GSSG Assay Kit (Promega GSH/GSSG Assay Kit) in a 96-well white plate. Cellular extracts were mixed with assay reagents to convert GSH and GSSG into a detectable luminescent product. Luminescence was measured using a plate reader, with the intensity proportional to the concentration of GSSG or total glutathione (GSSG+GSH). The data were normalized to total protein content to control for differences in sample concentrations. The GSH to GSSG ratio was then calculated to assess cellular oxidative status.

#### Detection of lipid peroxides by BODIPY™ 581/591 C11

The day before treatment 1.7 × 10^6^ SU-DHL10 cell were seeded in a 6-well plate in 2 mL of RPMI. Lipid peroxidation levels were performed by flow cytometry, using C11-BODIPY (581/591) C11 dye (**Table S4**). Cells were grown in a 6-well plate and incubated with 1.5 μM C11-BODIPY (581/591) for 20 minutes. Cells were then washed twice in HBSS and resuspended in 500 µL of HBSS. Lipid peroxide levels were determined in 50,000 cells at 581/591 nm (Ex/Em) excitation using the BD FACS SORP Aria Fusion flow cytometer. 100 μM was used as a positive control.

#### Oxidative stress

Reactive oxygen species (ROS) measurements were performed by flow cytometry (catalog in **Table S4**), using two different fluorescent dyes, Dihydroethidium (DHE, 2.5 mM) and MitoSOX (2 mM) to preferentially detect superoxide anion. Cells were grown in a 6-well plate and incubated with the dye for 30 min. Then, the cells were washed in PBS, resuspended in HBSS, and 50,000 cells were analyzed using a BD FACS SORP Aria Fusion flow cytometer using the following conditions: 518/606nm excitation/emission (Ex/Em) for DHE, at 396/610 nm Ex/Em for MitoSOX. We used cells treated for 1 h at 37 °C with 100 μM H_2_O_2_ as a positive control.

#### In vivo GCB PDX

Synergy studies were conducted using clinical-grade IKE (MedChem Express), and zotatifin, provided by eFFECTOR Therapeutics. Germinal Center B-cell-like (GCB) patient-derived xenograft (PDX) tumor (Public Repository of Xenografts, ProXe, DFBL-75549-R2) were established in male NOD scid gamma (NSG) mice aged over 8 weeks via surgical dorsal tumor implantation. Treatment described above (**Fig. 5A**, **Fig. S5C**).

#### In vivo A20

A20 cells expressing luciferase IVI were engrafted via tail vein injection (1 × 10⁶ cells per mouse) into BALB/c mice. See treatment above (**Fig. 6**). Zotatifin treatment was as indicated in results, and 1 × 10⁶ mouse CAR-T cells per mouse were given Day 5 via tail vein injection. The Region of Interest (ROI) was specified in the IVIS Spectrum Imaging Software to measure the total bioluminescent signal, or flux, from a particular area. The ROI was monitored twice weekly over a period of 43 days.

#### Quantification and Statistical Analysis

Proteomics analysis, RNA sequencing, and GSEA were conducted using RStudio (R 4.4.1, https://posit.co/download/rstudio-desktop/). GraphPad Prism (version 10.5, GraphPad Software) was used for all other analyses, including flow cytometry, ELISA, and viability assays. Specific test details are provided in the figure captions.

## Authors’ Disclosures

### Authors’ Contributions

P. Manara and J.H. Schatz design the project.

P. Manara, A. Newsam, VVG Saralamma and J.H. Schatz design CAR T experiments.

P. Manara, J.H. Schatz, D. Bilbao, M.V. Russo A. Newsam designed in vivo experiments. VVG Saralamma, A.B. Martinez, N Fattakhov performed in vivo experiments.

P. Manara, A.M. Barroso, A.M. Carbone, O.B. Lightfuss; D. Chahar performed bench assay.

K. Hoffman, P. Manara and T. A. Cunningham performed TMT-pSILAC

P. Manara, AYA, F. Maura performed bioinformatics.

P. Manara performed formal analysis.

P. Manara and J.H. Schatz wrote the first draft of the paper

All authors contributed to data interpretation and read, edited, and approved the manuscript

## Acknowledgments

We wish to thank Mr. Christian McDonald for Polysome Profiling and Dr. Anna Bianchi for technical assistance with CAR T Flow cytometry staining. We thank the laboratory of Naoki Hosen at Osaka University for sharing their *SLC3A2* knockout clone of U266 cells. Research in this publication was performed in part at the Cancer Modeling, Flow Cytometry, and Onco-Genomic Shared Resources (CMSR RRID: SCR022891, FCSR RRID: SCR022501, OGSR RRID: SCR022502) of the Sylvester Comprehensive Cancer Center at the University of Miami, supported by the National Cancer Institute (NCI) of the National Institutes of Health (NIH) under award P30CA240139. The content is solely the responsibility of the authors and does not necessarily represent official views of the NIH. We wish to thank Bioinformatics Solutions Inc for performing Mass Spectrometry.

