## Supplementary materials for "NRF2 translation block by inhibition of cap-dependent initiation sensitizes lymphoma cells to ferroptosis and CAR-T immunotherapy"


A. Supplementary Methods……………………………………………….1-2

B. Supplementary Figures with Legends.....…………………………...3-7

C. Supplementary Tables with Legends…………...……………….8-9

D. References……………………………………...………………. 10

### A. SUPPLEMENTARY METHODS

#### RNA-seq

We performed RNA sequencing (RNA-seq) on SU-DHL10 cells treated with zotatifin and compared them to DMSO controls. RNA was extracted from each sample and sequenced to an average depth of over 40 million raw paired-end 100 bp reads using the Illumina NovaSeq 6000 platform. Differential expression analysis was conducted with edgeR software, utilizing generalized linear models and the quasi-likelihood F-test. Genes were considered differentially expressed if they met the following criteria: false discovery rate (FDR) ≤ 0.05, logarithm of counts per million reads (logCPM) ≥ 0, and p-value ≤ 0.05. A log2 fold change (FC) ≥ ±1 was used to determine significant differential expression between the treated and control groups.

*GSEA*

Gene Set Enrichment Analysis (GSEA) was performed using the GSEA Desktop Application (https://www.gsea-msigdb.org/gsea/index.jsp) to identify enriched pathways. Differentially expressed genes were ranked by log2FC and used as input for the analysis. The analysis utilized the Hallmark gene set from the Molecular Signatures Database (MSigDB). Key parameters included 1,000 gene set permutations and the Signal2Noise metric for gene ranking. Pathways with an FDR q-value < 0.25 were considered significantly enriched. The results were visualized using the GSEA platform tools, and leading-edge genes contributing to significant pathways were further analyzed for their biological relevance.

*Comparison RNAseq vs TMT-pSILAC*

We compared RNA-seq and TMT-pSILAC data using R Studio by applying a logFC cutoff of ≥1 or ≤-1 to define significant changes, excluding genes with a P-value >0.05. We filtered both datasets to retain only genes present in both RNA-seq and TMT-pSILAC, then combined them for comparison. Bar plots were generated to visualize the logFC values for matched genes, illustrating expression patterns across the datasets and highlighting both consistent and divergent results.

#### TMT-pSILAC Enrichment

*GSEA*

For Gene Set Enrichment Analysis (GSEA) using the Hallmark gene sets, the fgsea package in RStudio was employed. Log2FC values from the TMT-pSILAC analysis were ranked and used to perform the GSEA. The Hallmark gene sets were sourced from a GMT file and loaded into the analysis. The fgsea function was executed with parameters including a minimum gene set size of 5, a maximum size of 1000, and 1000 permutations to evaluate the significance of enrichment. To visualize the results, a dot plot was created, showing the top enriched Hallmark pathways with gradient coloring based on the Normalized Enrichment Score (NES). The dot plot was saved as a PNG file for further presentation and analysis.

*Enrichr*

Enrichr analysis was conducted using the Enrichr platform (https://maayanlab.cloud/Enrichr) with the KEGG gene set library. Pathways with significant enrichment were identified based on A logFC cutoff of ≥ 1 or ≤ -1. The -log10(adjusted P-value) was calculated to assess pathway significance, with values ≤ -1 indicating upregulation and values ≥ 1 indicating downregulation."

#### EGFP NF-kB Reporter

The pHAGE-6x-NF-kB-conA-HygEGFP plasmid was a gift from D. Krappmann (Helmholtz Zentrum), used as previously described.(1) Infected cells were treated with 200ng/ml of doxycycline for 24 hours. Flow cytometry measured GFP as NF-kB readout compared to uninduced cells.

### B. SUPPLEMENTARY FIGURES WITH LEGENDS

### FIGURE S1

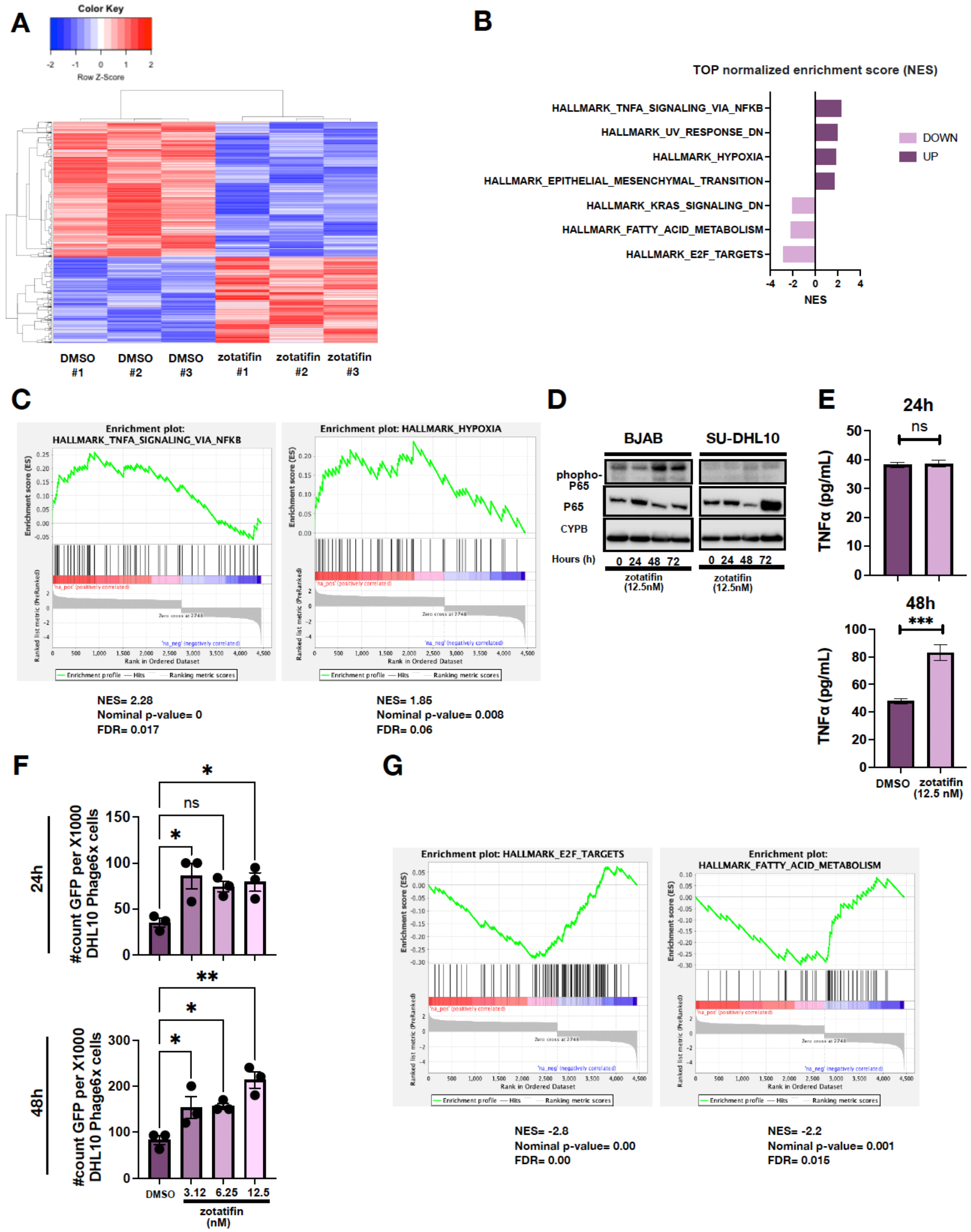

**Supplementary Figure S1 (preceding page):** A) RNA-seq heat map comparing Zotatifin (12.5 nM) vs. DMSO at 24 hours. Data form three independent experiment. B) GSEA Hallmark gene sets showing top upregulated and downregulated pathways with nominal p-value < 0.01 and FDR < 0.25. C) Enrichment plot of the top 2 upregulated pathways from Hallmark GSEA. D) Immunoblot depicting NFKB activation (pP65) in DLBCL cells (BJAB and SU-DHL10) treated with Zotatifin (12.5 nM). Data from two independent experiments. E) TNFα levels at 48 hours post Zotatifin (12.5 nM) treatment. Data represent mean ± SEM of three experiments. Statistical analysis was performed using repeated measures two-way ANOVA with Sidak's multiple comparison test (ns, ***P < 0.001). F) Histogram showing increased EGFP-positive cells in SU-DHL10 transfected with px458 (NFkB EGFP reporter) compared to untreated controls at 24 and 48 hours, upon various Zotatifin concentrations (3.125 nM, 6.25 nM, 12.5 nM). Statistical analysis was performed using one-way ANOVA with Bonferroni's multiple comparison test. Data represent the mean ± SEM of three independent experiments (ns, *P < 0.05, **P < 0.01 vs. DMSO). G) Enrichment plot of the top 2 downregulated pathways from Hallmark GSEA.

**Supplementary Figure S2 (next page):**
A) Immunoblot time course showing OCI-Ly3 (left) and SU-DHL10 (right) cells for SLC3A2 and puromycin, treated with 12.5 nM zotatifin for 0, 4, 6, 16, 24, and 48 hours. Data are from two independent experiments. B) TMT-pSILAC analysis of downregulated proteins with $\log_2$FC ≤-1. C) GSEA Hallmark analysis for the top 10 downregulated pathways from TMT-pSILAC data. D-E) Comparison of TMT-pSILAC upregulated (D) and downregulated (E) proteins with RNA-seq differentially expressed genes, where upregulation is defined as $\log_2$FC ≥1 and downregulation as $\log_2$FC ≤-1.

**FIGURE S2**

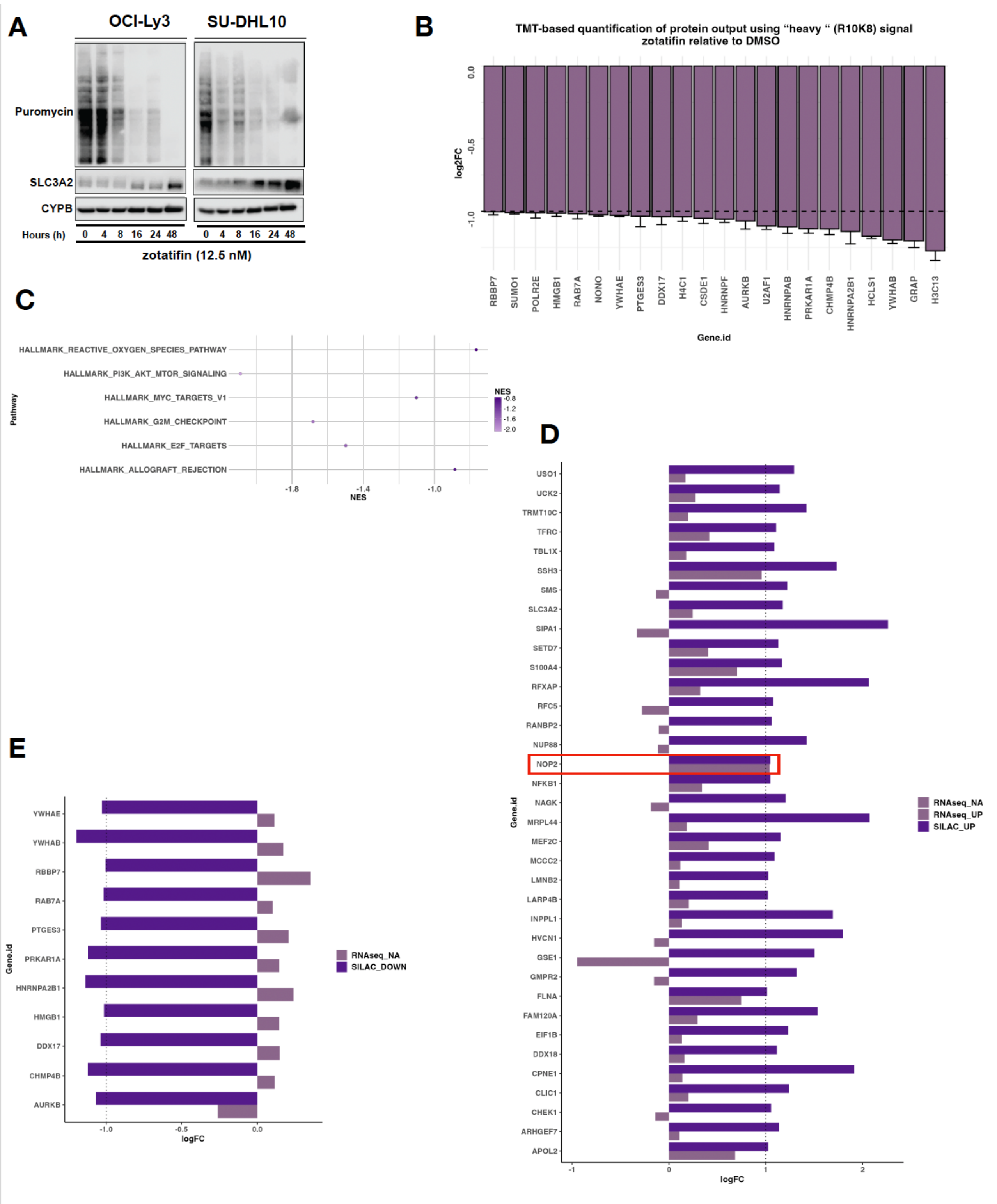
A
OCI-Ly3
SU-DHL10
Puromycin
SLC3A2
CYPB
Hours (h)
0 4 8 16 24 48
0 4 8 16 24 48
zotatifin (12.5 nM)
B
TMT-based quantification of protein output using "heavy " (R10K8) signal
zotatifin relative to DMSO
log2FC
0.0
-0.5
-1.0
RBBP7
SUMO1
POLR2E
HMGB1
RAB7A
NONO
YWHAE
PTGES3
DDX17
H4C1
CSDE1
HNRNPF
AURKB
U2AF1
HNRNPAB
PRKAR1A
CHMP4B
HNRNPA2B1
HCLS1
YWHAB
GRAP
H3C13
Gene.id
C
HALLMARK_REACTIVE_OXYGEN_SPECIES_PATHWAY
HALLMARK_PI3K_AKT_MTOR_SIGNALING
HALLMARK_MYC_TARGETS_V1
HALLMARK_G2M_CHECKPOINT
HALLMARK_E2F_TARGETS
HALLMARK_ALLOGRAFT_REJECTION
Pathway
-1.8
-1.4
-1.0
NES
NES
-0.8
-1.2
-1.6
-2.0
D
USO1
UCK2
TRMT10C
TFRC
TBL1X
SSH3
SMS
SLC3A2
SIPA1
SETD7
S100A4
RFXAP
RFC5
RANBP2
NUP88
NOP2
NFKB1
NAGK
MRPL44
MEF2C
MCCC2
LMNB2
LARP4B
INPPL1
HVCN1
GSE1
GMPR2
FLNA
FAM120A
EIF1B
DDX18
CPNE1
CLIC1
CHEK1
ARHGEF7
APOL2
Gene.id
-1
0
1
2
logFC
RNAseq_NA
RNAseq_UP
SILAC_UP
E
YWHAE
YWHAB
RBBP7
RAB7A
PTGES3
PRKAR1A
HNRNPA2B1
HMGB1
DDX17
CHMP4B
AURKB
Gene.id
-1.0
-0.5
0.0
logFC
RNAseq_NA
SILAC_DOWN

**Supplementary Figure S3:**

A) GSH/GSSG ratio in SU-DHL10 cells treated with zotatifin alone or ferroptosis inducer SASP (100nM). Statistical analysis: one-way ANOVA with Bonferroni multiple comparison test. Data represent mean ± SEM of three independent experiments. Statistical analysis used one-way ANOVA with Bonferroni multiple comparison test (ns, ****P < 0.0001). B) Immunoblot for cleaved caspase 3, XIAP, MYC, and GPX4 in DHL10 cells treated with DMF (50 μM)-zotatifin (12.5nM) at 24h. Data from two independent experiments.

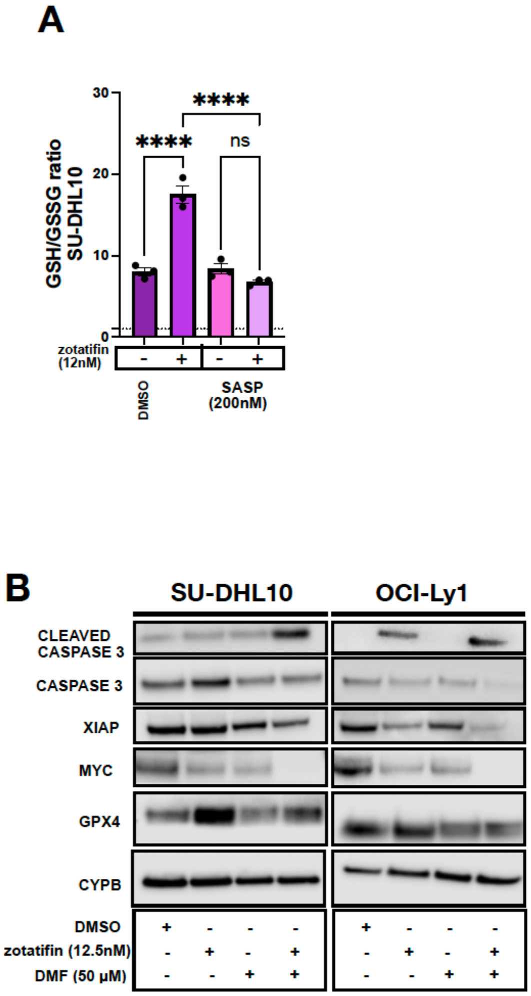

**Supplementary Figure S4.** A) Linear regression slope analysis of Cancer Cell Line Encyclopedia (CCLE) data for mRNA expression of HMOX1, NFE2L and KEAP1 versus CR-1-31B EC50, generated using the Dependency Map (DepMap) portal. C) Lipid peroxidation (left) and ROS measurement (right) with BODIPY C11 and MitoSOX in HAP1 cells WT (left) and F163L (right) treated with zotatifin and DMF. Data represent mean ± SEM of three independent experiments. Statistical analysis was performed using repeated measures two-way ANOVA with Sidak's multiple comparison test (***P < 0.001, ****P < 0.0001).

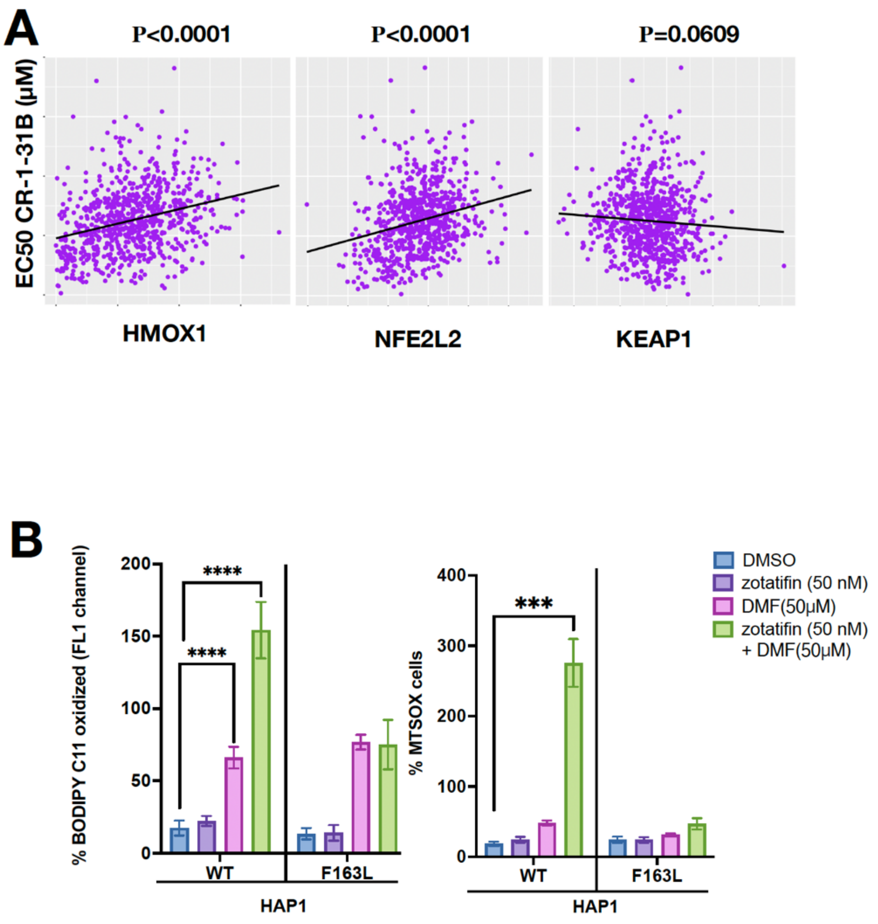

**Supplementary Figure S5.**

A) 8h co-culture of 4-1BB-CD19 CAR T cells with SU-DHL10 (left) and OCI-Ly1 (right), CFSE+ pretreated DLBCL (DMSO or 12.5 nM zotatifin). Measured by flow cytometry (see schematic 6C). B) 8h co-culture of CD28-CD19 CAR T cells (see schematic 6C) with SU-DHL10 (left) and OCI-Ly1 (right) cells for MitoSox to measure ROS and then fixed, permeabilized, and probed with CD3. Data represent mean ± SEM of three experiments. Statistical analysis was performed using repeated measures two-way ANOVA with Sidak's multiple comparison test (*P < 0.05, **P < 0.01). C) IFN γ levels for DHL10 treated with DMSO, Zotatifin for 32 hours (24h+8h) and Zotatifin(24h) washed for the last 8h. Data represent mean ± SEM of three experiments. Statistical analysis was performed using one-way ANOVA with Bonferroni's multiple comparison test. Data represent the mean ± SEM of three independent experiments (**P < 0.01 vs. DMSO). D) Average daily body weight ±SEM of surviving animals. Statistical analysis was performed using repeated measures two-way ANOVA with Sidak's multiple comparison test. E) Average total flux signal (p/s) over time for mice implanted with luciferase-expressing A20 cells. Represented logarithmic scale for the first eight days of treatments. Data represent mean ± SEM of ten independent experiments. Statistical analysis was performed using repeated measures two-way ANOVA with Sidak's multiple comparison test.

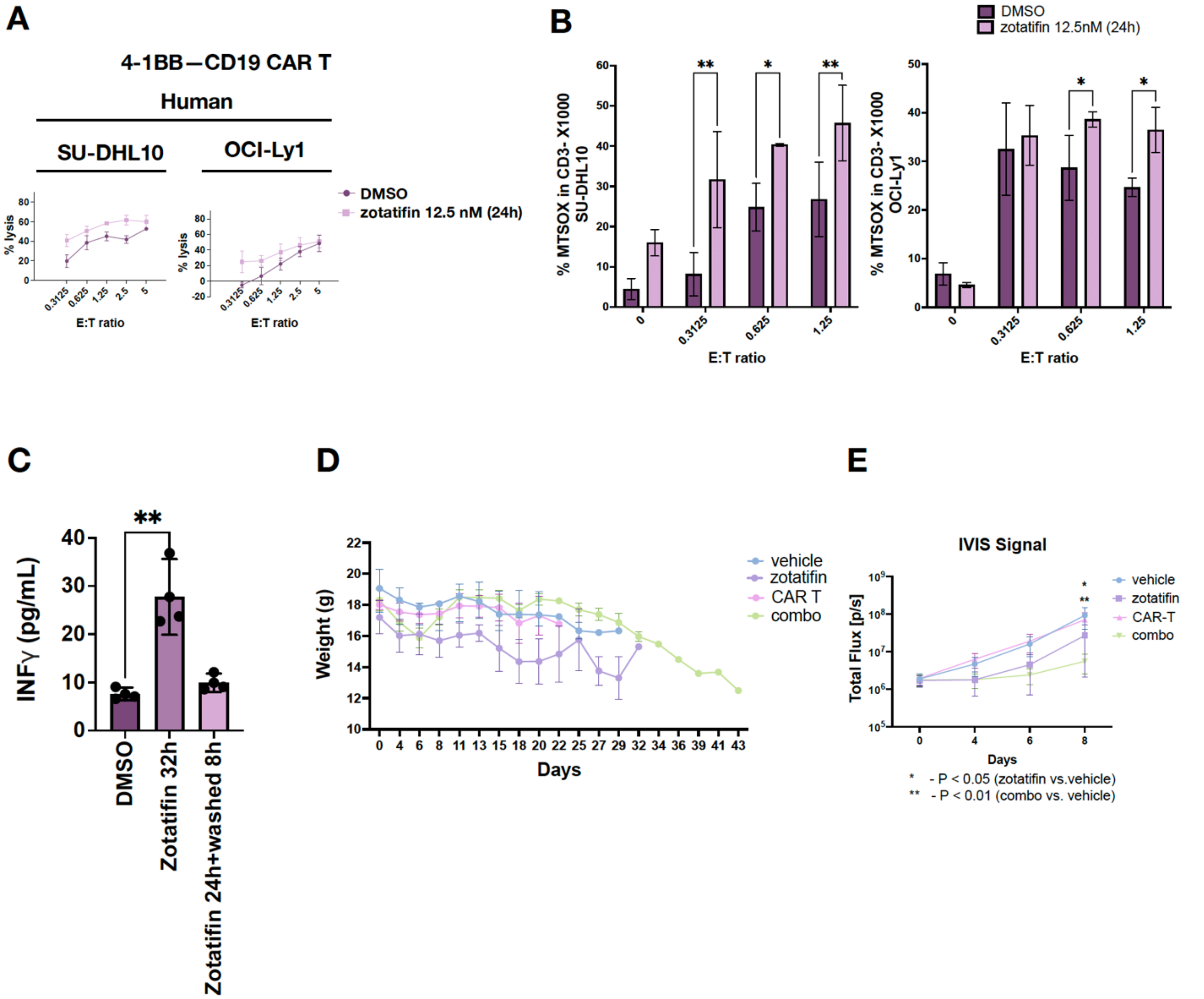

### C. SUPPLEMENTARY TABLES/LEGENDS

#### Supplementary Table S1: Reagents

| Compound | Company | Catalog # |
|---|---|---|
| Erastin | MedChemExpress | HY-15763 |
| RSL3 | MedChemExpress | HY-100218A |
| Dimethyl fumarate | MedChemExpress | HY-17363 |
| N-Acetylcysteine | MedChemExpress | HY-B0215 |
| Cycloheximide | MedChemExpress | HY-12320 |
| Altretamine | MedChemExpress | HY-B0181 |
| IFN-gamma Protein, Human | MedChemExpress | HY-P7025 |
| Zotatifin | MedChemExpress | HY-112163 |
| Imidazole Ketone Erastin (IKE) | MedChemExpress | HY-114481 |
| Sulfasalazine | Selleckchem | S1576 |
| Puromycin | Thermo Scientific | CHM01U363 |
| G418 (Geneticin) | InvivoGene | |

#### Supplementary Table S2: Western blotting antibodies

**Primary:**

| Marker | Species | Company | Catalog # |
|---|---|---|---|
| NRF2 (D1Z9C) | Rabbit anti-Human | Cell Signaling Technology | 12721S |
| XIAP (D2Z8W) | Rabbit anti-Human | Cell Signaling Technology | 14334S |
| GPX4 | Rabbit anti-Human | Cell Signaling Technology | 52455S |
| FTH1 | Rabbit anti-Human | Cell Signaling Technology | 3998S |
| HO-1 (E3F4S) | Rabbit anti-Human | Cell Signaling Technology | 43966S |
| CBS (D8F2P) | Rabbit anti-Human | Cell Signaling Technology | 14782S |
| 4F2hc/SLC3A2 (D6O3P) | Rabbit anti-Human | Cell Signaling Technology | 13180S |
| Cleaved Caspase-3 (Asp175) (5A1E) | Rabbit anti-Human | Cell Signaling Technology | 9664S |
| NF-κB p65 (D14E12) | Rabbit anti-Human | Cell Signaling Technology | 8242S |
| Phospho-NF-κB p65 (Ser536) (93H1) | Rabbit anti-Human | Cell Signaling Technology | 3033S |
| Anti-c-Myc [Y69] | Rabbit anti-Human | Abcam | ab32072 |
| NFS1 | Rabbit anti-Human | Proteintech | 15370-1-AP |
| CYPB | Rabbit anti-Human | ThermoFisher Scientific | PA1-027A |
| Anti-Puromycin | Mouse anti-Human | Sigma-Aldrich | MABE343 |
| β-Actin (8H10D10) | Mouse anti-Human | Cell Signaling Technology | 3700S |
| Anti-rabbit IgG, HRP-linked | Goat anti-Rabbit | Cell Signaling Technology | 7074S |

| | | | |
|---|---|---|---|
| Anti-mouse IgG, HRP-linked | Horse anti-Mouse | Cell Signaling Technology | 7076S |

**Primary housekeeping:**

| Marker | Species | Company | Catalog # |
|---|---|---|---|
| GAPDH (D16H11) | Rabbit anti-Human | Fluidigm | 5174S |
| CYPB | Rabbit anti-Human | ThermoFisher Scientific | PA1-027A |
| β-Actin (8H10D10) | Mouse anti-Human | Cell Signaling Technology | 3700S |

**Secondary:**

| Marker | Species | Company | Catalog # |
|---|---|---|---|
| Anti-rabbit IgG, HRP-linked | Goat anti-Rabbit | Cell Signaling Technology | 7074S |
| Anti-mouse IgG, HRP-linked | Horse anti-Mouse | Cell Signaling Technology | 7076S |

**Supplementary Table S3: qPCR primers**

| primers | Company | Catalog # |
|---|---|---|
| NFE2L2 | Quiagen | PPH06070A-200 |
| SLC3A2 | Quiagen | PPH00829A-200 |

18 sequences:
Forward: 5'- CAGCCACCCGAGATTGAGCA
Reverse: 5' - TAGTAGCGACGGGCGGTGTG

**Supplementary Table S4. Flow staining**

| staining | Company | Catalog # |
|---|---|---|
| CellTrace™ CFSE | ThermoFisher Scientific | C34554 |
| Anti-HumanCD98hc Monoclonal (5E5) | eBioscience | 11-0982-42 |
| Anti-Human IRF1 | BD Pharmingen | 566322 |
| BD Horizon™ BUV661 Mouse Anti-Human CD3 | BD Bioscience | 612964 |
| LIVE/DEAD™ Fixable Blue | Invitrogen | L23105 |
| BODIPY™ 581/591 C11 | Thermofisher Scientific | D3861 |
| MitoSOX™ Mitochondrial Superoxide Indicators | Invitrogen | M36008 |
| Dihydroethidium (DHE) | Invitrogen | D11347 |
